# MERRA-based trend analysis identifies complex, multidecadal patterns of changing climatic suitability for Cassin’s Sparrow (*Peucaea cassinii*)

**DOI:** 10.1101/2023.09.16.558057

**Authors:** John L. Schnase, Mark L. Carroll, Paul M. Montesano, Virginia A. Seamster

**Affiliations:** NASA Goddard Space Flight Center, Greenbelt, Maryland, 20708 USA; ADNET Systems, Inc., Bethesda, Maryland, 20817 USA; New Mexico Department of Game and Fish, Santa Fe, New Mexico, 87507 USA

## Abstract

Cassin’s Sparrow (*Peucaea cassinii*) is a grassland resident of the American Southwest. Despite decades of study, there remains uncertainty regarding the conservation status of the species. The species’ past response to a changing climate may help explain this uncertainty, especially if patterns vary across the species’ range. In this study, we combine data from NASA’s Modern-Era Reanalysis for Research and Applications, Version 2 (MERRA-2; M2) with field observations spanning the past 40 years to examine historical changes in climatic suitability for Cassin’s Sparrow across its full annual range within the continental United States. We examine two time- and variable-specific time series using MaxEnt. The M2 times series uses a mix of 30 microclimatic variables and ecosystem functional attributes related to energy and water fluxes for predictors; the MERRAclim (MC) time series uses 19 MERRA-2-derived bioclimatic variables for predictors. Trend analysis reveals complex patterns of slowly increasing climatic suitability over 69.5% of the study area in the MC time series accompanied by decreases over 24.4% of the area. Shifts in the study area-wide, weighted centroid for climatic suitability show a northwesterly, 40-year displacement of 1.85 km/yr. The M2 time series points to a less favorable history with increasing and decreasing trends over 54.9% and 40.1% of the study area, respectively, and a westerly weighted centroid shift of 2.60 km/yr. A clear subset of seven M2 and MC variables emerged as the most important determinants of suitability over the past 40 years. These variables also demonstrated complex patterns of non-constant trends across the study area. Suitability trends in both time series appear to have little in common with current, state-level, abundance-derived conservation status assessments. Increasing winds, drying land surface conditions, and variability in North American Monsoon rainfall appear to be dominating, climate-related influences on the species. We thus see complex patterns of historical change in climatic suitability depending on whether time series models are driven by bioclimatic variables alone or by variables more aligned with ecological function. This leads us to conclude that modeled estimates of climatic suitability for Cassin’s Sparrow can vary widely depending on the temporal frame, spatial extent, and environmental drivers considered, and that this variability mirrors the uncertainty in the literature regarding the species’ conservation status. Furthermore, these factors should be taken into account in future conservation assessments for the species, and retrospective ecological niche modeling, as applied here, offers a promising approach to addressing these issues.

## Introduction

Cassin’s Sparrow (*Peucaea cassinii* Woodhouse, 1852) is an elusive, ground-dwelling endemic of the arid grasslands of the southwestern United States (U.S.) and northern Mexico [1–3]. Like many grassland birds, evidence gathered over the past several decades suggests that Cassin’s Sparrow is experiencing a contraction of viable habitat and declining regional populations [4–8]. This is reflected in documents such as the State Wildlife Action Plan for New Mexico, which lists Cassin’s Sparrow as a declining species, susceptible to shifting environmental conditions or disease outbreaks that could lead to rapid population changes [9]. However, other sources report the species as stable [10–13]. Partners in Flight identifies Cassin’s Sparrow as a species of low conservation concern [14], and, of nine grassland birds in New Mexico, recent work has shown Cassin’s Sparrow to be the only species for which gains in suitable habitat are projected over the next 50 years [15]. NatureServe’s assessment is more complex, reporting Cassin’s Sparrow to be imperiled in Oklahoma, vulnerable in Kansas and Nebraska, and apparently secure in Texas, New Mexico, Arizona, and Colorado, while assigning the species a global ranking of demonstrably secure [16].

These differing views likely arise, in part, from natural history traits that complicate the observational record [3,5,17,18]. Cassin’s Sparrow is a grassland-shrub specialist [19]. A vegetation mix of grasses and low shrubs appears to be critical to the bird’s breeding ecology; however, considerable variation in the proportions of that mix seems to be well tolerated [2,3,5,19]. Cassin’s Sparrow also appears to be highly sensitive to precipitation, following the movement of monsoon rains throughout the breeding season in an itinerant pattern that is difficult to track or interpret [3,5,20]. Sorting out key ecological details, such as these, has been complicated by the species’ ground-dwelling habit. Often described as secretive, this non-descript sparrow lives much of its life on the ground, nearly invisible to even the most experienced observer, until the breeding season when males perform a flight display [2,3,5,21]. Males use this uncommon skylarking behavior to establish breeding territories, attract females, and maintain pair bonds. Launching from an exposed perch atop a low shrub, birds fly 10 m or more into the air, then flutter their wings in a slow descent to the ground or nearby perch while producing their distinctive, primary song [2].

A better understanding of the temporal dynamics of Cassin’s Sparrow’s historical response to a changing climate could help explain some of the uncertainties regarding the species’ conservation status and aspects of the species’ natural history and ecology that may be confounding the issue. We now understand that the multidimensional niche space of a species may shift, through adaptation or acclimation, as may its demography in response to climate changes [22]. We also know that climate change can drive shifts in the spatial distribution of many bird species as they track suitable conditions. These shifts are the consequence of multiple processes, including changes in habitat and the distribution of resources, dispersal patterns, and carrying capacity [23]. While climate-based ecological niche modeling (ENM) is commonly used to project anticipated responses by a species to a changing climate, less consideration has been given to retrospective patterns of change that could, in some cases, shed light on the present and future status of a species [24].

In this study, we combine data from NASA’s Modern-Era Reanalysis for Research and Applications, Version 2 (MERRA-2; M2) with field observations spanning the past 40 years to perform a retrospective ENM analysis of Cassińs Sparrow’s evolving environmental niche. In doing so, we assume that a combined look at historical patterns of change in suitable conditions alongside historical trends in the environmental drivers of those changes will provide new insights into the ambiguities surrounding Cassin’s Sparrow’s status. Our approach focuses on changes in climatic suitability and embodies a novel integration of three distinctive elements.

First, we base our analysis on environmental variables obtained solely from the M2 reanalysis. Climate reanalyses combine past observations with numerical models to generate a consistent time series of hundreds of fundamental, physical drivers of the Earth system. They offer a comprehensive description of Earth’s observed climate as it has evolved over the past half century at a fine temporal scale [25,26]. M2’s information-rich, long-running, high temporal-resolution data address ENM’s growing requirement for ecologically-relevant environmental predictors that can be tailored to specific modeling objectives while taking into account a species’ biological and ecological requirements [27–33].

The current study spans the 40-year period from 1980 to 2019 and is based on two climatic variable time series. In one, we use 30 M2 variables selected to reflect key attributes of the microclimate and climate-related ecosystem functioning [34,35]. The collection contains the precursor temperature and precipitation variables from which the 19 classic, bioclimatic predictors (i.e., bioclim variables) commonly used in ENM are derived [36,37]. In addition, it includes environmental attributes of more direct biological significance that are not explicitly represented in the 19 bioclim variables, such as soil moisture and evaporation from land, wind direction and speed, and various solar radiation fluxes [36,38–42]. In the second time series, we use 19 M2-derived bioclimatic variables modeled after the classic bioclim predictors [37,41]. Taken together, these two sets of predictors provide a more detailed view of macro- and microclimatic factors influencing environmental suitability for different species than could be realized by using the traditional bioclimatic predictors alone.

Our work with M2 variables complements a trend toward increased use of satellite-derived ecosystem functional attributes (EFAs) to improve model performance in ENM-based conservation practice [30,43–50]. In contrast to other types of predictors, EFAs, such as seasonal heat dynamics, the net energy fluxes that drive trophic webs, primary productivity, vegetation greenness, evapotranspiration, and soil moisture, relate to the performance of an ecosystem as a whole, and their values are potentially the consequence of multiple ecosystem processes [49]. ENMs that explicitly incorporate EFAs enable a more integrative perspective on environmental dynamics and allow for a more detailed characterization of the microclimatic conditions experienced by a species [49,51–53]. As a source for ENM predictors in our work, climate reanalyses provide a unique, readily-accessible mix of classic climatic variables and EFAs.

A second feature of the study is that we employ time-specific ENM in our analysis: our dependent and independent variables are temporally aligned across the time-span of the study. Detailed spatiotemporal information about the geographic distribution and changing dynamics of climatic suitability for a species is critical to conservation planning [30,43–46,48,50,54–56]. ENM is often applied within a time-averaged framework in which the values of environmental variables are averaged over time spans that are not in temporal registration with the occurrence records upon which models are calibrated or tested [55,57]. While useful for exploring species distributions at a broad level, modeling within a time-averaged framework can elide complex effects of the environment on an organism, especially highly mobile or behaviorally complex species [55,58–61]. Of particular concern to conservation work, studies have shown that temporal mismatches in the time period spanned by occurrence data and the climate baseline can decrease the utility and accuracy of ENM products [62–67].

In the current work, we use averaged values for our environmental variables across a sequence of eight, five-year time intervals spanning the 40-year period of the study. We then use time-specific species observations corresponding to these intervals as dependent variables in the models that form the basis of our analyses, thereby enabling a fine-grain, temporally-explicit view into historical patterns of changing climatic suitability.

Finally, we employ variable-specific ENM in our analysis: a tailored set of independent variables is used for each of the five-year intervals in the time series. There is an increasing awareness of the importance of variable selection in modeling environmental spaces [27,28,68–72]. At the same time, the increased availability of large and sometimes novel environmental data sets has made it difficult to select relevant predictors by anything other than automated or semiautomated means [27,29,69,70,73]. The third way the work described here is distinctive is in our use of NASA’s MERRA/Max system to do automatic variable screening within each of the temporally-aligned, five-year spans of our analysis.

MERRA/Max provides a scalable feature selection approach that enables direct use of global climate model (GCM) outputs in ENM [74,75]. The system accomplishes this selection through a MaxEnt-enabled Monte Carlo optimization that screens a collection of variables for potential predictors of suitable conditions. Based on a machine learning approach to maximum entropy modeling, MaxEnt is one of the most commonly-used software packages in the ENM modeling community [76–78]. Among its many features, MaxEnt ranks the contribution of predictor variables to the formation of its models. MERRA/Max’s Monte Carlo method exploits this capability in an ensemble strategy whereby many independent MaxEnt runs, each drawing on a random pair of variables from within a large collection of variables, converge on a global estimate of the top contributing variables in the collection being screened. Importantly, the ensemble’s bivariate MaxEnt runs can operate in parallel in a high-performance, cluster computer environment, and, with a sufficient number of processors available, can do so in only a few minutes, regardless of variable collection size.

With MERRA/Max, variable selection is guided by the indirect biological influences injected into the algorithm’s selection process by the species occurrence files, identifying biologically and ecologically plausible predictors in large, multidimensional data sets where selection through ecological reasoning or other means is not feasible [69,74,79]. In the current study, the time-specific variable selection performed by MERRA/Max enables a view into the changing patterns of environmental determinants of climatic suitability that would otherwise be difficult, if not impossible, to obtain.

Collectively, these three aspects of the study offer a more detailed look at past patterns of change in the climatic suitability for Cassin’s Sparrow than have been previously reported. In the sections that follow, we describe our method and results, discuss what we see as the important take-away lessons, and conclude with recommendations for next steps.

## Materials and Methods

We obtained the data used in this study from publicly available sources. We developed scripts operating in the open-source R v4.1.3, Python 2.7.12, Java 1.8.0_261, and bash v5.0 environments to perform the computational work of the study. Input data and principal scripts used in the study are available for download at: https://github.com/jschnase/MMX_Toolkit [80].

### Occurrence data

To create a set of longitudinal occurrence records, we used the R rgbif library to obtain Cassin’s Sparrow observations from the Global Biodiversity Information Facility (GBIF) [81,82] (GBIF.org; [15 January 2022] GBIF Occurrence Download https://doi.org/10.15468/dl.x33grq). A total of 32,518, georeferenced records for the years 1980 through 2019 were downloaded. More than 95% of the records were originally sourced from the eBird citizen-scientist observational dataset [83], but the download also included research-grade records from iNaturalist [84], the National Ecological Observational Network’s (NEON) breeding land bird point count collection [85], and museum specimen records from the National Museum of Natural History (NMNH) [86], American Museum of Natural History (AMNH) [87], Harvard University’s Museum of Comparative Zoology [88], Kansas University [89], and the University of Arizona [90].

We merged the observations into a time-series comprising eight, five-year aggregated collections: 1980–84, 1985–89, 1990–94, 1995–99, 2000–2004, 2005–09, 2010–2014, and 2015-19 (Fig 1A). These collections ranged in size from 263 records in the 1980 group to over 14,000 records in the 2015 collection. To reduce sampling bias, we applied two filtering steps. To avoid the potential of double counting the same individual bird, we thinned the entire dataset to non-overlapping observations within a 16 km (∼10 mile) buffer based on the species’ home range size [2,5,17,21]. For count uniformity across the series and to reduce record-density influences that can diminish model performance [91–94], we obtained random, 250-record samples for each five-year span in the time series, which proved in subsequent tuning steps to be an optimal sample size in the final models.

**Fig 1.**
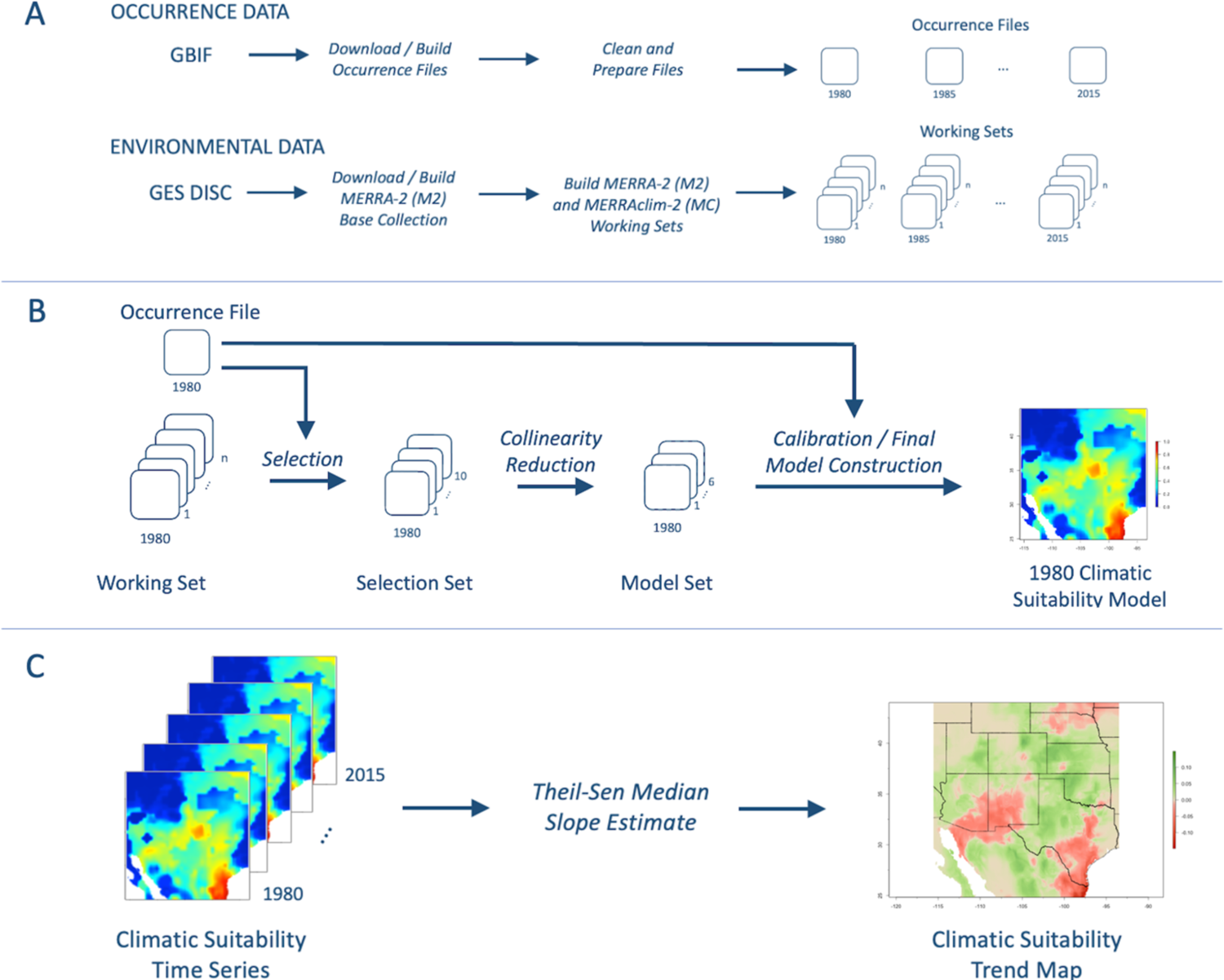
Study workflow. The major processing steps used in the study include (A) data preparation, (B) time series construction, and (C) time series analysis. Data sources include the Global Biodiversity Information Facility (GBIF) and the Goddard Earth Sciences Data and Information Services Center (GES DISC).

### Environmental variables

We used data from the U.S. Geological Survey (USGS) National Gap Analysis Program (GAP) to define a study area that encompasses the species’ range across the continental U.S. (latitude 24.8°N to 44.0°N and longitude 93.5°W to 115.6°W) [95]. We then created a *base collection* of 30 M2 variables that we judged from personal experience and knowledge of the literature to be potentially important environmental determinants of suitable conditions for Cassin’s Sparrow (Table 1, Fig 1A) [2,3,5,17,18]. These were drawn from four, hourly, time-averaged, two-dimensional collections in which each variable represented one surface-level spatial grid across the landscape: (1) M2T1NXSLV, consisting of air temperatures, wind components, total precipitable water vapor, etc., at popularly-used vertical levels, (2) M2T1NXFLX, consisting of surface fluxes, such as observation-corrected total precipitation, surface air temperature, specific humidity, wind speed, and re-evaporation, (3) M2T1XRAD, consisting of radiation estimates, such a surface albedo, cloud area fraction, cloud optical thickness, solar radiation, and (4) M2T1NXLND, which is made up of an assortment of variables of particular interest to environmental suitability modeling applications, such as surface soil wetness, root zone soil wetness, soil temperatures at various layers, and important elements of the land energy and water balance equations [34,96].

**Table 1.**
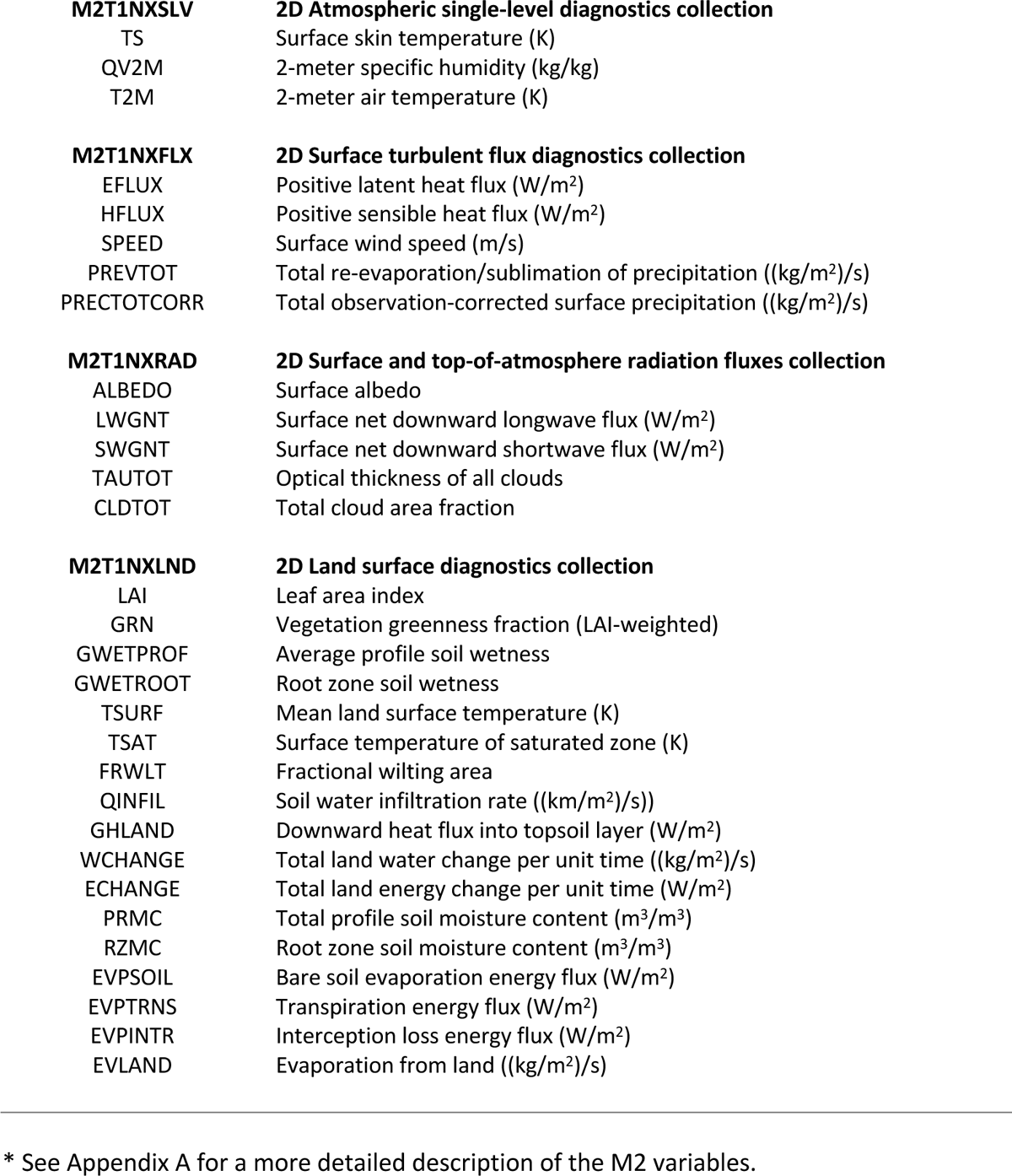
MERRA-2 (M2) variables.*

We obtained M2 data in its native, Network Common Data Form 4 (NetCDF4) format from research collections housed in the NASA Center for Climate Simulation (NCCS) [97]; however, the data are also available to the general public through the Goddard Earth Sciences Data and Information Services Center (GES DISC) [98,99]. We used the subsetting capabilities of Python’s xarray v2022.06.0 library [100] to assemble global, five-year aggregate collections from the original downloads. These groupings corresponded to the eight intervals of our occurrence data time series. Each five-year collection contained the averaged maximum, minimum, and mean values for the 30 selected variables.

From this base collection, we assembled eight *working set* collections of averaged mean values for the 30 variables, which we tailored to the specific requirements of the study using R’s rgdal v1.5-18 library [101]. Each environmental layer was clipped to the spatial extent of the study area, which encompasses the geographic range of Cassin’s Sparrow, re-projected, and formatted for use by MaxEnt following the method of Hijmans et al. [102]. The resulting eight working set collections corresponded to the eight intervals of our occurrence data time series. To smooth the representation of local environmental conditions, we resampled the M2 layers from their native spatial resolution of 1/2 ° latitude × 5/8 ° longitude to 5.0 arc-minutes (1/12 °) resolution (∼7.6 km at latitude 35.0°N, which is within the study area) using the R raster v3.6-3 [103] library’s bilinear interpolation routine.

In addition to the M2 collections, we built base and working set collections of M2-derived bioclimatic variables, which we refer to as the MERRAclim-2 (MC) collections (Table 2). MC’s variables are modeled on Worldclim’s classic 19 bioclim variables, which were designed to highlight climate conditions generally understood to relate to a species’ physiology [36,37]. The bioclim variables are derived from monthly maximum and minimum temperature values and the average values for monthly precipitation. We built the M2 version of the bioclim dataset using the R dismo v1.3-9 library’s ‘biovars’ function [104] following the method used by Vega et al. to generate the research community’s first generation MERRAclim collection [105]. In their original formulation, Vega et al. [105] used the first-generation MERRA’s two-meter temperature (T2M) and two-meter specific humidity (QV2M) variables as inputs. In an improvement over the earlier version of MERRA, the current, updated MERRA-2 provides a modeled, observation-corrected total precipitation (PRECTOTCORR) variable expressed as a mm/sec rate, which we adapted for use in MC by converting to a monthly amount [106].

**Table 2.** MERRAclim-2 (MC) variables.

|  |  |
| --- | --- |
| MC_Bio01 | Annual mean temperature (°C) |
| MC_Bio02 | Mean diurnal temperature range (°C) |
| MC_Bio03 | Isothermality [(MC_Bio02/MC_Bio07)*100] (%) |
| MC_Bio04 | Temperature seasonality [(Standard deviation*100)] (°C) |
| MC_Bio05 | Maximum temperature of the warmest month (°C) |
| MC_Bio06 | Minimum temperature of the coldest month (°C) |
| MC_Bio07 | Temperature annual range (MC_Bio05-MC_Bio06) (°C) |
| MC_Bio08 | Mean temperature of the wettest quarter (°C) |
| MC_Bio09 | Mean temperature of the driest quarter (°C) |
| MC_Bio10 | Mean temperature of the warmest quarter (°C) |
| MC_Bio11 | Mean temperature of the coldest quarter (°C) |
| MC_Bio12 | Annual precipitation (mm) |
| MC_Bio13 | Precipitation of the wettest month (mm) |
| MC_Bio14 | Precipitation of the driest month (mm) |
| MC_Bio15 | Precipitation seasonality (Coefficient of variation) (%) |
| MC_Bio16 | Precipitation of the wettest quarter (mm) |
| MC_Bio17 | Precipitation of the driest quarter (mm) |
| MC_Bio18 | Precipitation of the warmest quarter (mm) |
| MC_Bio19 | Precipitation of the coldest quarter (mm) |

### Time series construction

Using the GBIF occurrence dataset, we constructed two time series spanning the years 1980 to 2019 in eight, five-year intervals. In the first, we used the M2 collections of working set variables; in the second, we used the MC working sets. A two-step processing workflow was applied to each five-year interval of each time series (Fig 1B). First, we used MERRA/Max to screen the working set variables in each of the eight, five-year spans of the 40-year series. We ran MERRA/Max on a dedicated testbed comprising a set of ten, 10-core Debian Linux 9 Stretch virtual machines (VMs) in the NCCS’s Explore high-performance cluster computing environment [107]. Using MERRA/Max’s standard screening configuration [74] and a per-variable sampling rate of 100, we performed three screening runs, each returning a set of top ten selected variables. We averaged the results of the three runs to create a *selection set* of the top ten overall predictors for each five-year interval. We then used variance inflation factor (VIF) analysis to reduce collinearities in the selected predictors. VIF shows the degree to which standard errors are inflated due to the level of multicollinearities [108]. We calculated the VIF for the selected variables using the R usdm v1.1-18 library and removed the least contributing variable in any pair of variables having r > 0.8, r^2^ > 0.8, and VIF > 10.0 [108,109]. This resulted in a final, *model set* of top-recommended predictors for each five-year interval in the two time series that had few if any multicollinearity issues.

Next, in a model calibration and final model construction step, we used the R ENMeval v2.0.3 package [110,111] and MaxEnt v3.4.4 [112] to identify optimal feature class (FC) and regularization multiplier (RM) settings for each set of occurrences and predictors in each of the five-year intervals of the M2 and MC time series. MaxEnt’s FCs are mathematical transformations of the predictors that allow complex environmental dependencies to be modeled and include linear (L), quadratic (Q), product (P), threshold (T), and hinge (H) settings; the RM controls how closely-fitted the output distributions will be [76,113]. We tuned parameters by performing a series of model runs across all possible combinations of six FCs (L, LQ, H, LQH, LQHP, and QHP) and eight RM values ranging from 0.5 to 4.0 in increments of 0.5. The combination of settings resulting in the lowest value for Akaike’s information criterion corrected for small sample size (AICc) [114] was taken to be an optimal tuning configuration, which was subsequently used to construct a final model for each five-year interval in the two time series [110,111]. For each of the 48 ENMeval calibration runs, and in the final MaxEnt model run, we used 10,000 background locations randomly selected from across the study area and performed a 10-fold cross validation in which 70% of the occurrences were selected for training and 30% for testing in each repetition [115,116]. Cloglog output scaling was used throughout. The cloglog format gives an estimate between 0.0 and 1.0 of the probability of presence, which, in the current study, we use as a proxy for environmental suitability [77]. The model calibration and final model construction step was performed in triplicate for each five-year interval to produce an averaged result that we used in the subsequent time series analysis.

### Time series analysis

We adapted the approach of Stephens et al. [117] to assess the rate of change in Cassin’s Sparrow climatic suitability between 1980 and 2019. We used the Theil-Sen method to create summary maps of the study area for each time series that showed regions of positive and negative trend in suitability over the 40-year span of the study (Fig 1C). The Theil-Sen median slope estimator provides a non-parametric means of robustly fitting a line to a set of points by finding the median slope of all the lines through all pairs of points in the set. It is relatively insensitive to outlying points and can be significantly more accurate than using simple, least squares linear regression. In our case, the Theil-Sen median slope estimate was applied across the eight raster layers comprising the final models of climatic suitability for each five-year interval of our two 40-year time series. There were, thus, eight points in the slope calculation for each cell in the resulting summary trend maps. We used the Mann-Kendall test to determine the statistical significance of the resulting trends [118–120]. R’s spatialEco v2.0-0 library was used for the Theil-Sen and Mann-Kendall analyses [121].

We used the movement of weighted centroids to characterize the direction and speed of shifts in climatic suitability. Weighted centroids represent the geometric center of all pixels in the studied area weighted by their suitability index [122]. We chose centroids for this purpose because of their ability to account for information derived across a species’ entire modeled range [123–125]. Our determination of the direction and speed of centroid movement was made by measuring the vector between weighted centroids of the 1980 and 2015 probability maps using R’s enmSdmX library [126]. The direction of shifts was quantified in degrees ranging between 0° and 360°, with 0° being due north.

We complemented our look at biological responses with an analysis of trends in the environmental variables underlying the time series models. First, we examined trends in the values of the most contributory variables across the entire time series. Top contributing variables were those that appeared in half or more of the 24 final models in the triplicated time series; these variables were ranked by their permutation importance, which can range from zero (no contribution) to one (high importance) [69,127]. For these study-wide top contributing variables, we computed Theil-Sen slopes to assess trends and visualize patterns of change in the mean values of each of M2’s and MC’s modeled variables over the past 40 years across the study area. We then examined trends in the relative contributions of each of these top variables over time (i.e., trends in each variable’s permutation importance) on a per-interval basis across the 40-year span of the study.

The final models produced in the calibration and final model construction step were judged for reasonableness based on first-hand knowledge of the species and its climatic preferences, what is known about Cassin’s Sparrow’s range from the published literature [2,3,5,17], and observational records from Cornell Lab’s eBird database [128]. To gain a quantitative perspective on performance, we used the Area Under the Receiver Operating Characteristic (ROC) Curve (AUC) as a measure of model accuracy. AUC is a measure of a model’s ability to discriminate presence from absence and ranges from 0.0 to 1.0 [129,130]. In addition, we used the True Skill Statistic (TSS) [131,132] and Percent Correctly Classified (PCC) [133] as measures of model accuracy, where higher values in both cases also indicate greater accuracy.

## Results

### Model performance

In evaluating model performance, we regarded AUC measures < 0.7 as low, 0.7–0.9 as moderate, and > 0.9 as high; similarly, we regarded TSS and PCC measures < 0.5 as poor, 0.5–0.8 as useful, and > 0.8 as good [134]. By these metrics, models exhibited moderate performance across all 24 models per time series in their mean AUC, PCC, and TSS values (Table 3). We judged this level of performance to be a reasonable basis for examining the broad trend patterns displayed by the time series models, which were the primary focus of the study [131–133]. Our distribution maps corresponded well with what is known about the natural history of the species and maps derived from observational records [128].

**Table 3.** Model performance.*

| Time Series | AUC | PCC | TSS |
| --- | --- | --- | --- |
| MERRA-2 (M2) | $0.817 \pm 0.009$ | $0.718 \pm 0.023$ | $0.528 \pm 0.017$ |
| MERRAclim-2 (MC) | $0.812 \pm 0.007$ | $0.734 \pm 0.008$ | $0.789 \pm 0.020$ |

\* Overall model performance for the M2 and MC time series. Mean ± standard error used throughout, $n=24$ in both time series.

### Climatic suitability

The study area encompasses approximately 3.91 x 10^6^ km^2^. Final models in the two time series reveal changing patterns of estimated climatic suitability for Cassin’s Sparrow across the region over the past 40 years that reflect differences in the type of predictors used and, presumably, actual changes in climatic suitability itself. On visual assessment, the most favorable climatic conditions for the species have generally concentrated in the southeastern regions of the study area according to both the M2 and MC time series; however, over time, a northwesterly shift in areas of high suitability is apparent in both series (Figs 2A, 3A). The Theil-Sen trend maps also show a northwesterly movement in increasing climatic suitability over the past 40 years, although the patterns of change are more broadly diffuse in the M2 series, as described in greater detail below (Figs 2B, 3B). An overall increase in climatic suitability was identified over approximately 54.9% of the study area (∼21.49 x 10^5^ km^2^) in the M2 time series and 69.5% of the study area (∼27.21 x 10^5^ km^2^) in the MC series (Table 4). Areas showing an overall decrease in climatic suitability ranged in size from approximately 40.1% (∼15.72 x 10^5^ km^2^) in the M2 time series to 24.4% of the study area (∼9.54 x 10^5^ km^2^) in the MC series.

**Fig 2.**
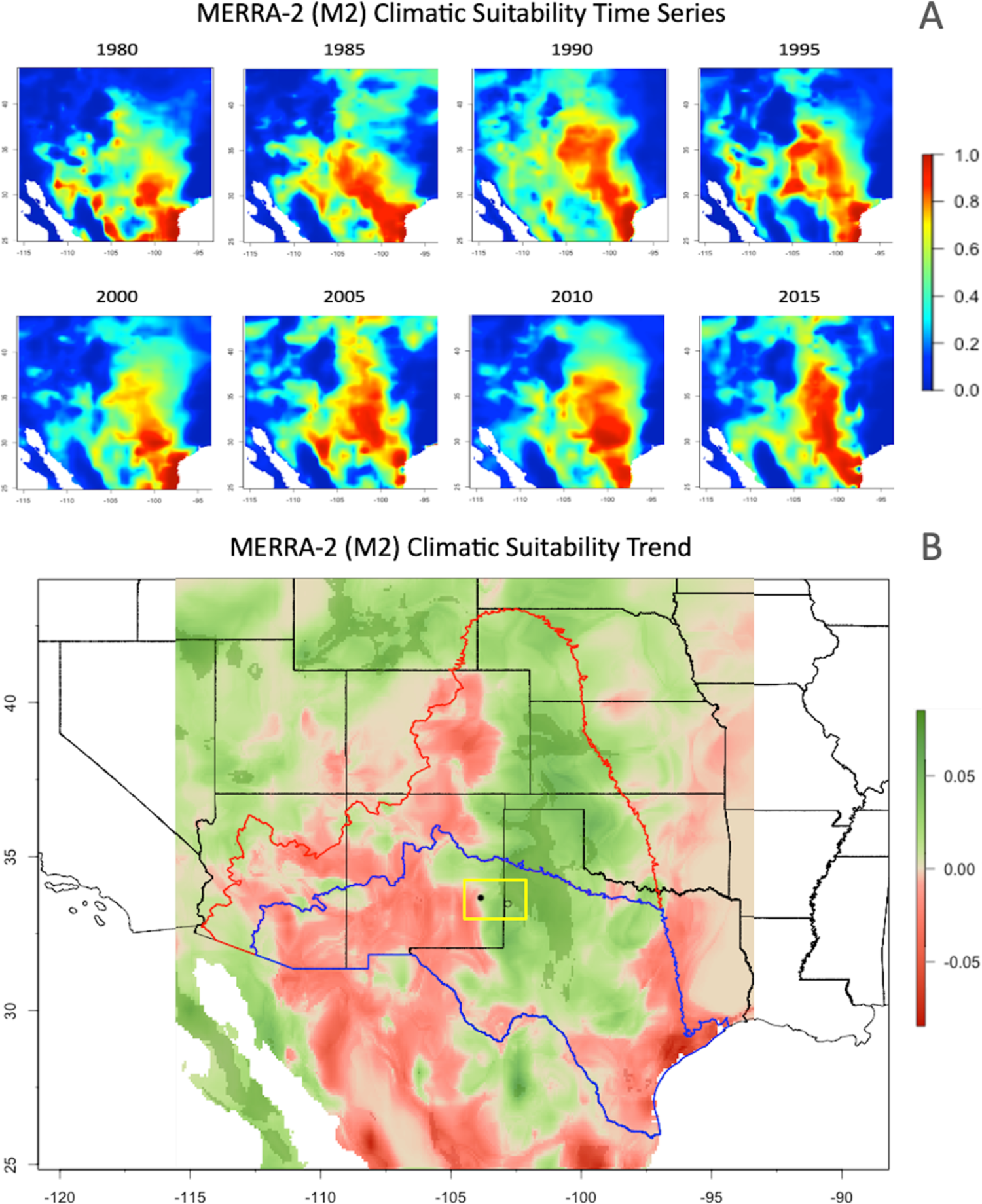
MERRA-2 (M2) climatic suitability trends. Averaged results from the three M2 time series runs: (A) Estimated probabilities of climatic suitability for Cassin’s Sparrow for the five-year intervals spanning 1980 to 2019. Probability values range from 0.0 to 1.0 with warmer colors indicating more favorable conditions. (B) Spatial distribution of Theil-Sen slopes showing the rate of change in probabilities of climatic suitability per five-year interval across the 40-year time series. Positive trends are indicated in green, negative trends in red. Statistically-significant positive and negative trends at the 95% confidence level are shown in dark green and red, respectively. Colored outlines indicate the northern extent of Cassin’s Sparrow’s U.S. breeding (red) and non-breeding (blue) ranges. Cassin’s Sparrow’s summer, breeding range encompasses all of the species’ winter, non-breeding range. The yellow box shows the location of a shift in weighted centroids for climatic suitability from 1980 (○) to 2015 (●).

**Fig 3.**
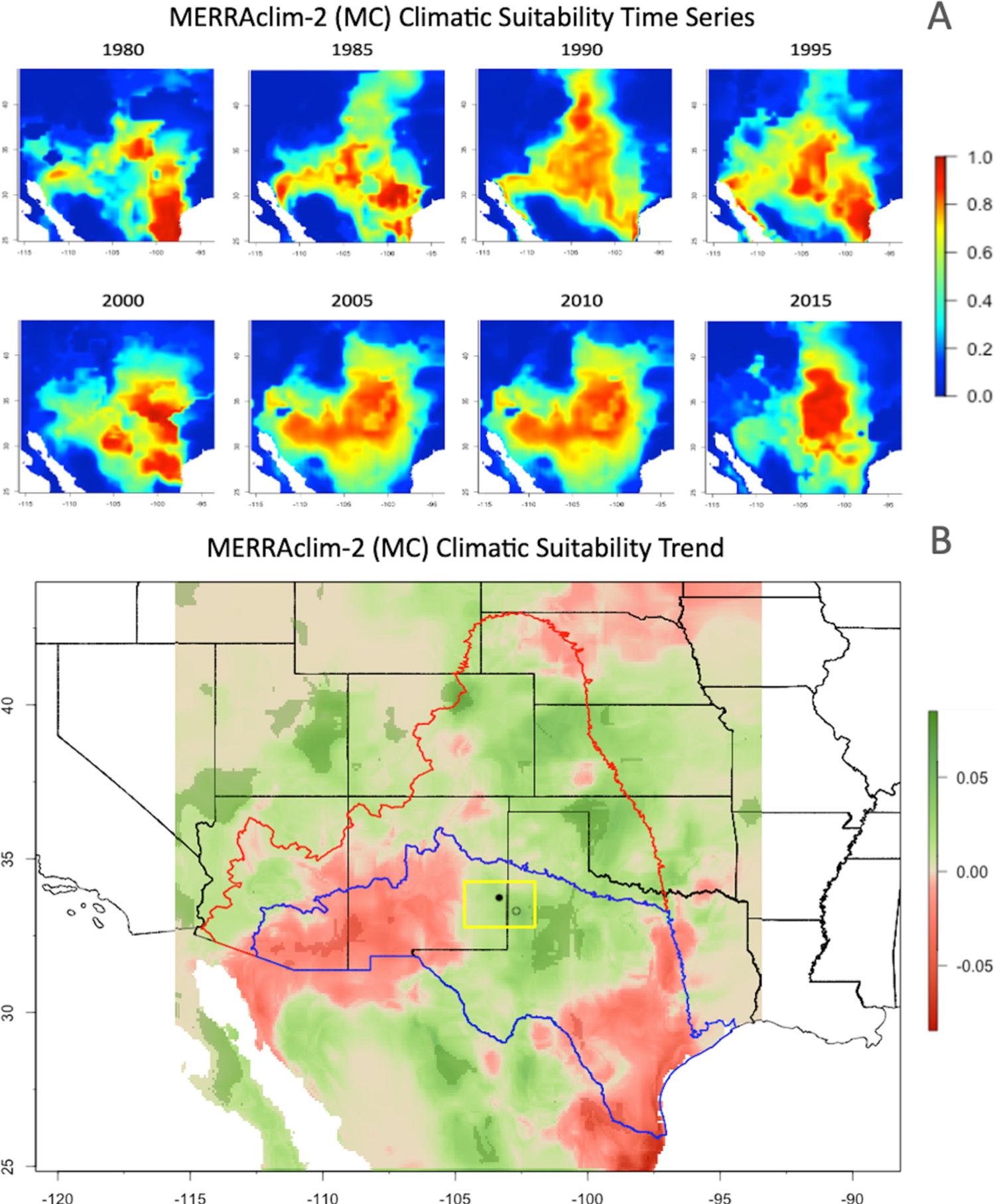
MERRAclim-2 (MC) climatic suitability trends. Averaged results from the three MC time series runs. See Fig 2 caption for additional detail. Theil-Sen results for the MC time series paint a similar picture. Statistically-significant probability estimates of suitable climate have been increasing by an average of 0.03 every five years for the past 40 years across 5.5% of the study area, with an accompanying average decrease in estimated probabilities of 0.06 over 0.9% of the region (Table 4). Significant positive trends concentrated in three clusters, one each in the northwest and northeast, the third in a central southwestern region, which, by contrast, showed a negative trend in the M2 analysis.

**Table 4.** Climatic suitability trends.*

| Time Series | Overall Trends |  | Significant Trends |  |  |  |
| --- | --- | --- | --- | --- | --- | --- |
|  | km<sup>2</sup> x 10<sup>5</sup> | % of area | Z | km<sup>2</sup> x 10<sup>5</sup> | % of area | TS Δ / 5-yr |
| **MERRA-2 (M2)** | 21.49 ± 3.96 | 54.9 ± 10.0 | 3.09 ± 0.01 | 2.41 ± 0.64 | 6.2 ± 1.6 | 0.03 ± 0.05 |
|  | 15.72 ± 1.28 | 40.1 ± 3.2 | -2.85 ± 0.15 | 0.19 ± 0.07 | 0.5 ± 0.2 | -0.05 ± 0.01 |
| **MERRAclim-2 (MC)** | 27.21 ± 1.07 | 69.5 ± 2.6 | 3.34 ± 0.08 | 2.15 ± 0.11 | 5.5 ± 0.3 | 0.03 ± 0.01 |
|  | 9.54 ± 0.06 | 24.4 ± 2.7 | -2.09 ± 0.01 | 0.37 ± 0.05 | 0.9 ± 0.1 | -0.06 ± 0.02 |

\* Table presents a summary of results obtained from Theil-Sen median slope analysis of the study's two time series. Trend areas are calculated for the map regions in Figs 2B and 3B that exhibited change at any level of statistical significance (Overall Trends) as well as statistically-significant change at the 95% confidence level (Significant Trends). Results are based on the three trend maps produced by the triplicated runs in each time series and show mean ± standard error across these three maps. Positive (green) and negative (red) trends are indicated by the corresponding positive and negative Mann-Kendall Z scores. Theil-Sen slope values (TS) represent the rate of change in estimated probabilities of climatic suitability per five-year interval across the 40-year time series in areas showing statistically-significant trends.

Focusing on areas of statistically-significant change provides a somewhat different perspective. The absolute values of positive and negative change across the region are low, and the areas where change can be identified as significant at the 95% confidence level are relatively small (Table 4). While significant trends are hard to ascertain for time series in which n=8, for the M2 series, the statistically-significant Theil-Sen results indicate that the estimated probabilities of suitable climate have been increasing by an average of 0.03 every five years for the past 40 years across 6.2% of the study area, with a corresponding average decrease in estimated probabilities of 0.05 over 0.5% of the region. Significant positive trends were concentrated in the central, northwestern, and far southwestern regions of the study area, while negative trends concentrated in the southeast and central southwest, imparting a west to northwesterly axis to these positive shifts that is consistent with the weighted centroid analysis, as described below.

However, there was a significant negative trend for the southeast region of the MC series that coincides with the pattern observed for the M2 series. Overall shifts in the climatic suitability for Cassin’s Sparrow were observed in both time series. The 40-year displacement of the weighted centroid for suitability in the M2 time series was approximately 104 km and had a westerly inclination (282°). This pattern contrasted with that seen in the MC time series, which showed a 40-year displacement of 75 km along a northwesterly trajectory (309°) (Fig 4). Centroid movement in the M2 time series shows a higher change velocity than that seen in the MC time series (2.60 km/yr vs. 1.85 km/yr respectively).

**Fig 4.**
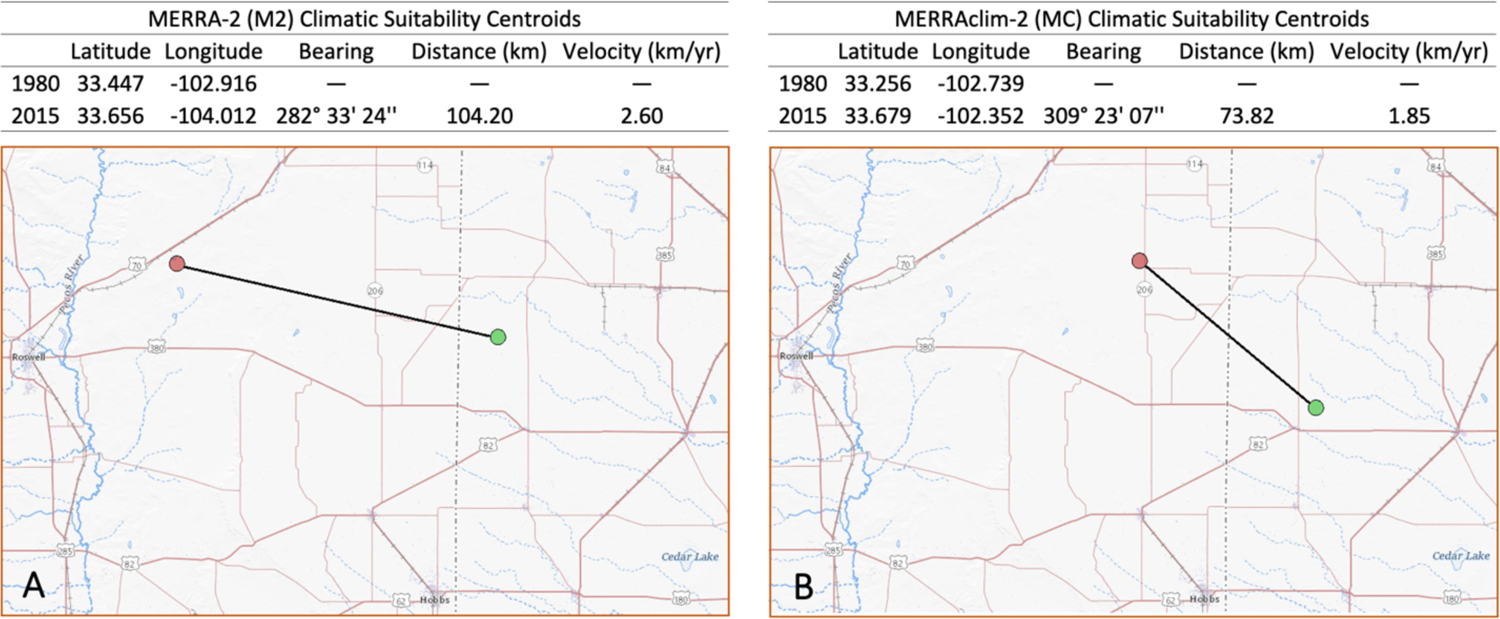
Climatic suitability shifts. Maps show direction, distance, and velocity of 40-year shifts in the weighted centroids for climatic suitability in (A) the M2 time series and (B) the MC time series from 1980 (●) to 2015 (●). For orientation, Roswell, NM is the city on the western boundary of the maps. Base map courtesy of the USGS with annotations by the authors.

### Environmental variables

The top contributing variables in the M2 time series were surface wind speed (SPEED), bare soil evaporation energy flux (EVPSOIL), 2-meter specific humidity (QV2M), interception loss energy flux (EVPINTR), optical thickness of all clouds (TAUTOT), total re-evaporation of precipitation (PREVTOT), and surface net downward shortwave flux (SWGNT) (Table 5).

**Table 5.** Top contributing variables.*

| MERRA-2 (M2) | n | PI | Description |
| --- | --- | --- | --- |
| SPEED | 24 | 18.3 ± 1.7 | Surface wind speed (m/s) |
| EVPSOIL | 24 | 12.4 ± 1.9 | Bare soil evaporation energy flux (W/m2) |
| QV2M | 24 | 6.2 ± 1.2 | 2-meter specific humidity (kg/kg) |
| EVPINTR | 20 | 19.3 ± 1.5 | Interception loss energy flux (W/m2) |
| TAUTOT | 17 | 9.9 ± 1.8 | Optical thickness of all clouds |
| PREVTOT | 15 | 16.4 ± 2.9 | Total re-evaporation/sublimation of precipitation ((kg/m2)/s) |
| SWGNT | 13 | 20.8 ± 2.8 | Surface net downward shortwave flux (W/m2) |

| MERRAclim-2 (MC) | n | PI | Description |
| --- | --- | --- | --- |
| MC_Bio08 | 24 | 39.6 ± 3.1 | Mean temperature of the wettest quarter (°C) |
| MC_Bio14 | 19 | 11.1 ± 1.0 | Precipitation of the driest month (mm) |
| MC_Bio18 | 18 | 23.2 ± 2.9 | Precipitation of the warmest quarter (mm) |
| MC_Bio03 | 16 | 10.7 ± 2.0 | Isothermality [(MC_Bio2/MC_Bio07)*100] (%) |
| MC_Bio15 | 15 | 10.6 ± 1.3 | Precipitation seasonality (Coefficient of variation) (%) |
| MC_Bio05 | 15 | 9.2 ± 3.7 | Maximum temperature of the warmest month (°C) |
| MC_Bio13 | 12 | 8.1 ± 2.6 | Precipitation of the wettest month (mm) |

\* Table presents a summary of the top contributing variables in the study's M2 (top) and MC (bottom) time series. Variables are ordered first by the number of times the variable appeared in the final models of the eight, five-year intervals of the triplicated times series (n), followed by the variable's mean permutation importance (PI) ± standard error across those appearances.

Theil-Sen analysis showed overall, positive trends across the study area for SPEED, QV2M, PREVTOT, and SWGNT, with positive change centrally located in the study area for SWGNT, concentrated in northern, northeastern, and southeastern regions in QV2M, and broadly scattered in SPEED and PREVTOT (Fig 5). Predominantly negative trends were seen in EVPSOIL and EVPINTR, with EVPSOIL’s negative changes concentrated in the southwest and EVPINTR’s broadly distributed across the entire study area. Nearly equivalent areas of overall positive and negative change were observed for TAUTOT, with positive change concentrated in the north and southeast (Table 6). M2 trends generally lacked statistical significance.

**Fig 5.**
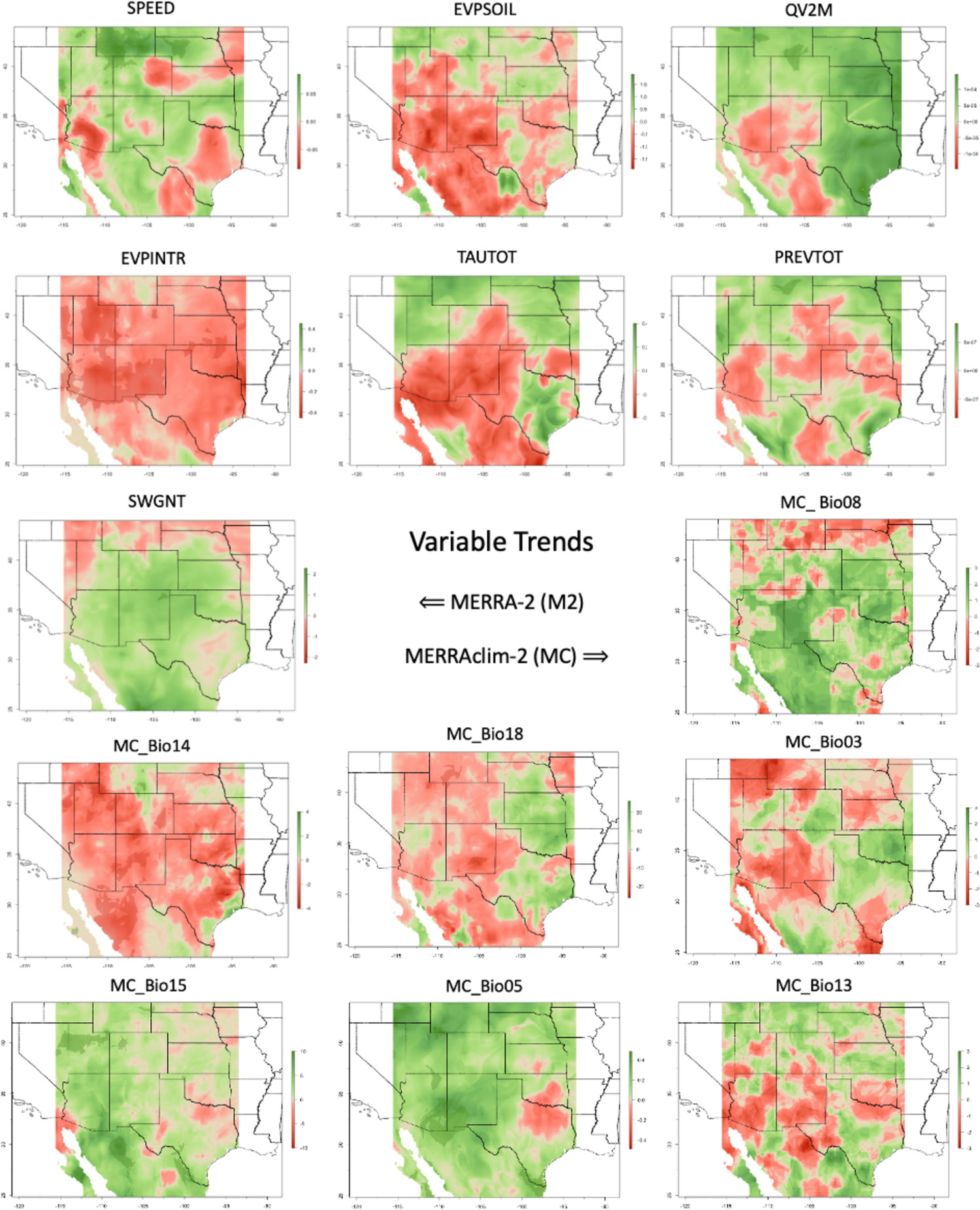
Variable trends. Maps show the Theil-Sen slope estimates for the top contributing variables in the M2 and MC time series (Table 5). Positive trends are shown in green; negative trends are shown in red. Color intensity represents the rate of change in the units of measure for the variable per five-year interval across the 40-year time series. Maps created by the authors.

**Table 6.**
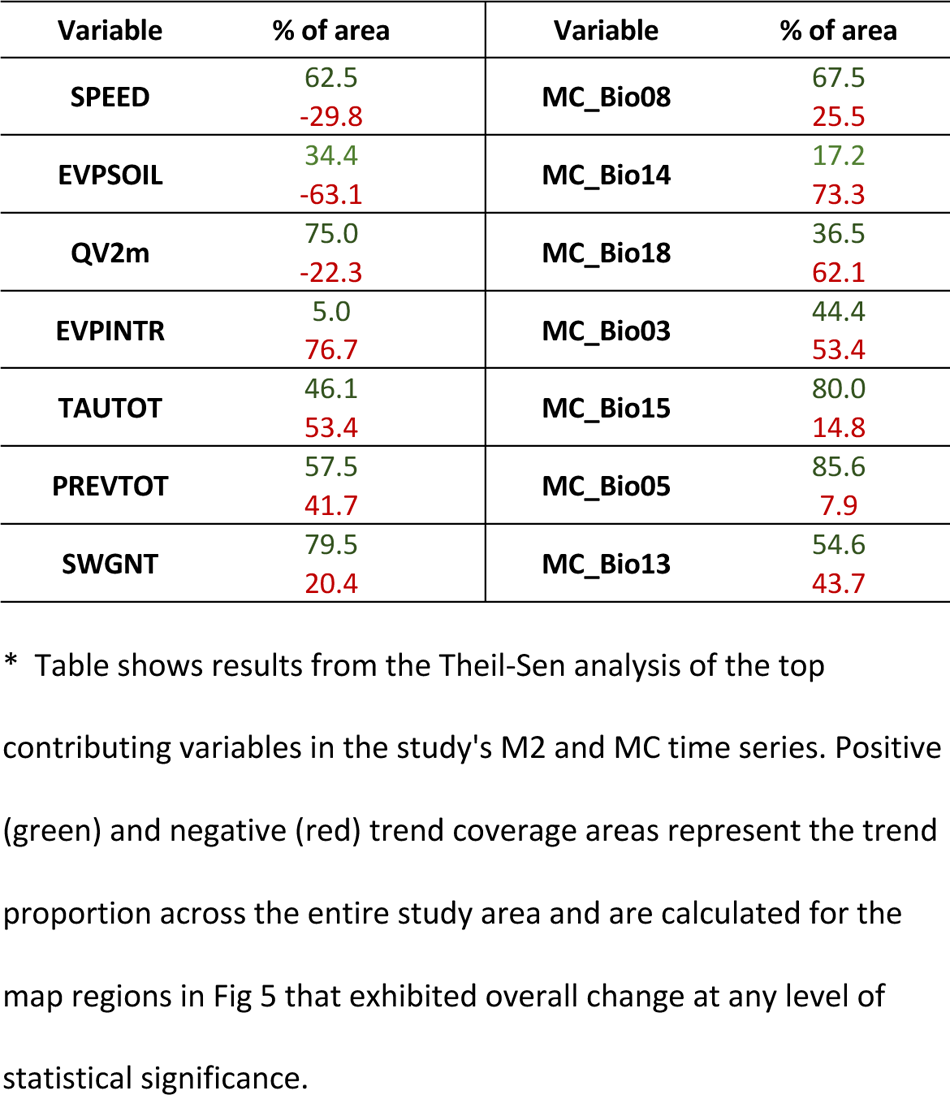
Variable area trends *.

| Variable | % of area | Variable | % of area |
| --- | --- | --- | --- |
| SPEED | 62.5<br>-29.8 | MC_Bio08 | 67.5<br>25.5 |
| EVPSOIL | 34.4<br>-63.1 | MC_Bio14 | 17.2<br>73.3 |
| QV2m | 75.0<br>-22.3 | MC_Bio18 | 36.5<br>62.1 |
| EVPINTR | 5.0<br>76.7 | MC_Bio03 | 44.4<br>53.4 |
| TAUTOT | 46.1<br>53.4 | MC_Bio15 | 80.0<br>14.8 |
| PREVTOT | 57.5<br>41.7 | MC_Bio05 | 85.6<br>7.9 |
| SWGNT | 79.5<br>20.4 | MC_Bio13 | 54.6<br>43.7 |

\* Table shows results from the Theil-Sen analysis of the top contributing variables in the study's M2 and MC time series. Positive (green) and negative (red) trend coverage areas represent the trend proportion across the entire study area and are calculated for the map regions in Fig 5 that exhibited overall change at any level of statistical significance.

The top contributing variables in the MC time series were mean temperature of the wettest quarter (MC_Bio08), precipitation of the driest month (MC_Bio14), precipitation of the warmest quarter (MC_Bio18), isothermality (MC_Bio03), precipitation seasonality (MC_Bio15), maximum temperature of the warmest month (MC_Bio05), and precipitation of the wettest month (MC_Bio13) (Table 5). The Theil-Sen studies showed predominantly positive trends across the study area for MC_Bio08, MC_Bio15, and MC_Bio05, with large expanses of positive change concentrated centrally and in the northwest region of the study area for MC_Bio05, concentrated in the southwest for MC_Bio08, the west and southwest for MC_Bio15 (Fig 5). Negative trends dominated in MC_Bio14 and MC_Bio18, with westerly-inclined, broadly diffuse patterns of change generally concentrated in the west for both variables. Nearly equivalent areas of positive and negative change were seen in MC_Bio03 and MC_Bio13, with broadly scattered patches of change apparent for each variable throughout the study area (Table 6). Most of the trends observed in the MC variables also lacked statistical significance.

Finally, time-specific variable selection also enabled a view into each variable’s contributory trend across the 40-year span of the time series (Fig 6). In the M2 time series, we observed a generally increasing trend in the relative contribution of EVPINTR to each five-year interval’s final model and a generally decreasing trend in the contributions of QV2M and SWGNT. SPEED and EVPSOIL were consistently high contributors across the board; TAUTOT and PREVTOT were consistently moderate contributors. In the MC time series, MC_Bio14 and MC_Bio03 show generally increasing trends; MC_Bio08 showed sharply decreasing trends. MC_Bio18 was a consistently high contributor across the series, with MC_Bio15, MC_Bio05, and MC_Bio13 making consistent contributions at moderate to low levels.

**Fig 6.**
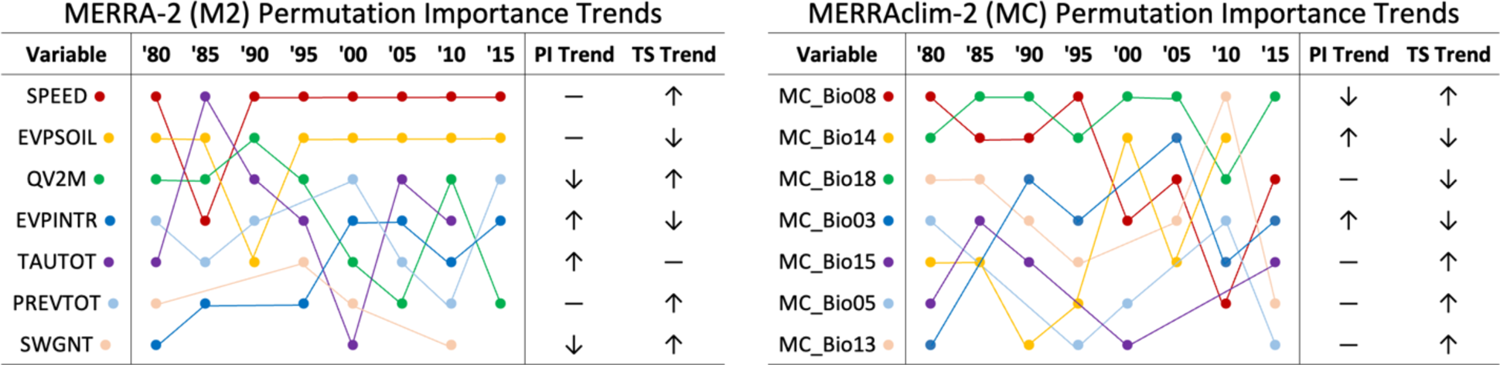
Permutation importance trends. Diagrams showing the relative contributions of the top contributing variables across the five-year intervals of the M2 (left) and MC (right) time series over the 40-year span of the study. The PI Trend column indicates whether a variable’s relative contribution to a five-year interval’s final model is generally increasing (↑), decreasing (↓), or remaining constant (–) across the 40-year span of the series. For comparison, the TS Trend column shows each variable’s overall net positive or negative Theil-Sen trend coverage area from Tables 6. Variables are listed in descending order of overall contribution to the time series, as shown in Table 5.

## Discussion

### Climatic suitability

The most important finding in the current study is that climatic suitability for Cassin’s Sparrow has been changing over the past 40 years, and those changes are manifest across the landscape as a complex pattern of positive and negative trends that can yield varying and sometimes contradictory interpretations depending on how suitability is modeled and the context in which modeled results are viewed. From a study-wide perspective, the northwesterly shift in favorable conditions we observed across both time series is consistent with what has been reported for Cassin’s Sparrow and many North American grassland bird species [8,135–137]. In addition, our velocity results are on par with other estimates. Bateman et al. [8], for example, found an average bioclimatic velocity of 1.27 km/yr to the west, northwest, and north over the past 60 years in the potential breeding distributions of 285 species of land birds across the continental U.S., with some potential breeding populations shifting at rates up to 5.51 km/yr.

Likewise, a recent study by Huang et al. [124] reported a mean bioclimatic velocity of 2.25 km/yr in 29 species of grassland birds along predominantly east-west axes of increasing environmental suitability, accompanied by an estimated, mean abundance-based velocity of 5.02 km/yr along more northerly inclined axes of increasing abundance. This leads us to believe we are seeing meaningful patterns with the approach we have taken. Assuming that is the case, these broad results belie potentially important details revealed by a more granular view.

For example, the MC time series paints a more favorable picture of changing conditions than what is seen in the M2 time series: climatic conditions are generally improving in the former, in the later we see an overall decline. This contrast is particularly apparent across the western regions of the study area and within the USGS GAP boundaries of Cassin’s Sparrow’s northernmost breeding range. In the west, M2-driven model results portray a northerly shift in improving conditions across the full extent of the study area. With the MC-driven models, changes are not nearly as distinct nor are they as extensive. M2-driven model results show sharply improving climatic suitability in the northeastern extent of the breeding range and a fall-off along the western boundaries of its breeding and non-breeding ranges, while the MC-driven model results show improving climatic suitability in the northeastern region and a fall-off along only the western boundary of Cassin’s Sparrow’s breeding range.

The differences between the two time series become even more noticeable when the scope is reduced to the state level. Seven states comprise Cassin’s Sparrow’s breeding and non-breeding ranges within the continental U.S.: Arizona (AZ), Colorado (CO), Kansas (KS), Nebraska (NE), New Mexico (NM), Oklahoma (OK) and Texas (TX) (Figs 2B, 3B; Table 7). According to the GAP range maps, Cassin’s Sparrow is found in four of these states only during the breeding season (i.e., CO, KS, NE, and OK). With KS and OK, we see similar trend patterns across the two time series. On the other hand, for CO, we see a sharp decline in suitability in the M2 time series, especially in the central part of the state, and a general improvement in conditions in the MC time series. We see the inverse pattern for NE across the two time series.

**Table 7.**
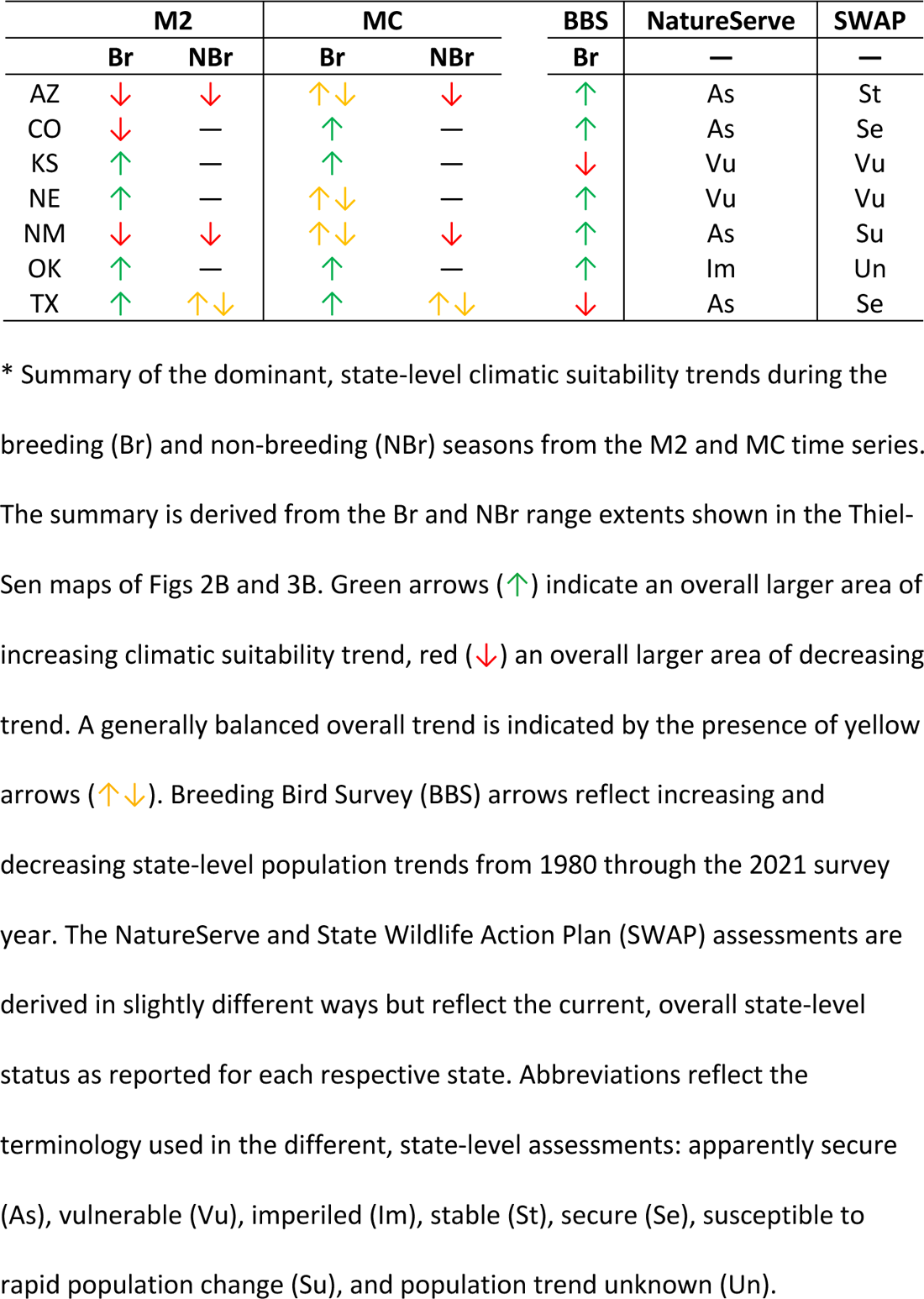
Comparison of state-level trends.

Within the U.S., Cassin’s Sparrow has historically been most abundant in AZ, NM, and TX [2,3,5,138,139]. These states represent the heart of Cassin’s Sparrow’s U.S. range, and it is here we see the species move from non-breeding to breeding areas between winter and summer months. This is also where we see the most striking differences in the M2 and MC time series’ model results. In the M2 series, climatic suitability in areas that historically accommodated seasonal range expansion, between the non-breeding and breeding seasons, is shown to have been declining over the past 40 years in AZ and NM. In contrast, climatic suitability in these areas appears to be improving or largely unchanged in the MC series. In TX, M2 model results indicate improving breeding season conditions in northern regions and decreasing suitability along the state’s southern border in breeding and non-breeding areas; similar but less pronounced patterns are observed in the MC time series.

Taken together, these findings are consistent with recent work showing that the nature, direction, and magnitude of changes in a species’ climatic niche are not uniform, but rather multidimensional in nature and shaped by complex, species-specific interactions that are sensitive to environmental drivers other than temperature and precipitation alone [122,124,140,141]. The findings also draw us into a consideration of the complexities that exist in the relationship between population abundance and environmental suitability that may speak to the ambiguities we see surrounding Cassin’s Sparrow’s conservation status.

Demographic attributes, such as abundance, are a crucial component of essentially all species conservation decision-making processes. For example, population size is a factor in NatureServe’s conservation status evaluations [142], which, together with the USGS’s North American Breeding Bird Survey (BBS) population trend estimates [143], contribute in some way to nearly all U.S. SWAPs and the identification of Species of Greatest Conservation Need (SGCN) [144]. Given that data on species occurrences are far more common and readily accessible than data on species abundance, it is not surprising that a significant amount of research has focused on clarifying the abundance-suitability (AS) relationship and the extent to which ENM-derived estimates of environmental suitability can serve as a proxy for population abundance [145]. While much remains to be learned about modeling the distribution of abundance, a consensus appears to be emerging around the following points:

1. the distribution of environmental suitability based on bioclimatic variables alone generally shows little, if any, correlation with the distribution of abundance [124,146–148];
2. the combination of bioclimatic predictors with other environmental attributes, such as EFAs, edaphic variables, topographic data, vegetation indices, etc., can capture the influence of factors affecting abundance rather than just occurrence, thereby yielding suitability model results that are often highly correlated with abundance [145,147–149], and, in some cases, reflect well the mean and maximal local abundances of a species [150];
3. the AS correlation is particularly strengthened when the added non-climatic predictors capture, in one way or another, environmental drivers that influence fitness, dispersal, recruitment, or other demographic properties of a species [149,151–153]; and
4. correlative ENM that takes into account this bioclimatic-demographic connection can provide practical benefits to spatial conservation efforts, such as readily-attainable, large-scale abundance estimates; hot-spot identification; and reduced survey and monitoring costs [24,145,147,148,150].

Our findings are consistent with the first point: we see different patterns of historical change in environmental suitability depending on whether time series models are driven by bioclimatic variables alone or by variables more aligned with ecological functioning, and therefore potentially species demography. We also see that state-level, climatic suitability trends in both time series appear to have little in common with three major, state-level, abundance-driven conservation status assessments in force today (Table 7). Finally, the differences we observe among the suitability distributions of both time series’ five-year interval maps (Figs 2A, 3A), point to varying and potentially important underlying temporal dynamics. This leads us to believe that modeled estimates of Cassin’s Sparrow’s distribution based on climatic suitability can vary widely depending on the temporal frame in which they are constructed, the spatial extent considered, and the environmental drivers being used. Furthermore, this variability can lead to differing conclusions regarding the conservation status of the species, which likely mirror the ambiguities we see in the current literature regarding the status of Cassin’s Sparrow. We cannot address the remaining points relating to the AS relationship at the moment, because we did not combine M2 and MC variables in the current study, nor did we directly compare our suitability results with demographic results. However, as described below, points (2) through (4) do help us frame what we believe are important next steps in this line of research.

### Environmental variables

It is not surprising that the two collections of independent variables used to drive our two time series of models should result in different patterns of historical climatic suitability change. Studies have consistently demonstrated that correlative suitability models based on bioclimatic variables alone capture different environmental dynamics compared to models based on attributes of potentially more direct biological significance to a species [27,49,154]. Further, while temperature and precipitation are far and away the most frequently used predictors in ENM studies, and often end up being the most important [155], it has been shown that conditions beyond these two variables exert a strong influence on bird distributions [8,156]. It is also not surprising that the MC-based time series shows more favorable historical trends overall than what we observed in the M2 time series. Studies have shown that the use of bioclimatic predictors alone tends to overestimate range size, presumably because of their inability to effectively account for all of a species’ use of space and all the microclimatic influences to which the species may be responding [157]. What we do find interesting is what appears to be the enhanced capacity to discriminate between different forcings on a species over time that results from combining time-specific variable selection with time-specific ENM.

In earlier work, we found evidence that MERRA/Max’s Monte Carlo approach to automated variable selection was capable of surfacing ecologically-plausible predictors from multivariate collections [74,75,158–160]. Given the developmental nature of the approach, that work focused on applying the technique to a single temporal frame of reference. Here, variable selection has been applied to specific time intervals across a 40-year time series. This gives us an opportunity to consider two types of trends in the selected environmental predictors: we can examine how the most influential determinants of climatic suitability change over time and the extent to which the relative contributions of those variables change over time.

Take, for example, the three most important variables in the MC time series: mean temperature of the wettest quarter (MC_Bio08), which appears to have been generally increasing across Cassin’s Sparrow’s range over the past 40 years while declining in importance as an environmental driver; total precipitation of the driest month (MC_Bio14), which has been decreasing across Cassin’s Sparrow’s range while increasing in importance as an environmental driver over the same period of time; and precipitation of the warmest quarter (MC_Bio18), which has also been generally decreasing across the species’ range, but across a discontinuous patchwork of positive and negative trends while remaining one of the top environmental drivers over the 40-year span of the study (Table 6, Fig 6). Collectively, these three variables appear to confirm that the long-running drought conditions across southwestern North America have had a major influence on climatic suitability for Cassin’s Sparrows [161,162]. That MC_Bio18 is consistently the most important contributing variable across the time series is particularly notable. A case can be made that MC_Bio18 essentially characterizes the seasonal precipitation trends of the North American Monsoon, which has a recognized influence on the life history and breeding biology of the species.

The North American Monsoon is a product of seasonal change in the atmospheric circulation patterns that occurs as the summer sun heats the continental land mass. During much of the year, the prevailing wind over northwestern Mexico, Arizona, and New Mexico is westerly and dry. As the summer heat builds over North America, a region of high pressure forms over the Southwest, and the wind becomes more southerly, drawing moisture from the Pacific Ocean, Gulf of California, and Gulf of Mexico. In the Continental U.S., this circulation brings thunderstorms and rainfall to Arizona, New Mexico, Southern California, Utah, and Colorado, providing much of the region’s annual total precipitation [163–167]. Importantly, apart from the spatial variability that is directly related to topography, there is much intraseasonal and interannual variability in the intensity and extent of monsoon rainfall, particularly in the southwestern U.S. and northwestern Mexico [168], which may explain the broken and overall weaker trend patterns observed in our results for MC_Bio18 (Fig 6).

Cassin’s Sparrow’s responsiveness to rainfall and apparent ability to quickly relocate throughout the summer months in order to track optimal conditions for breeding has been a repeating theme in the literature for nearly a century: it is a life history trait that has been the source of much uncertainty about the species’ range and abundance [2,3,5,18,20,139]. The model contribution trends of the three top predictors in the MC time series, when set alongside landscape-scale trends in the variables themselves, appear to point to the importance of monsoon precipitation over Cassin’s Sparrow’s southwestern North American range over the past 40 years as temperatures and overall drying conditions have increased across the region [161,162]. The next two most contributory variables are isothermality (MC_Bio03) and precipitation seasonality (MC_Bio15). Isothermality represents temperature evenness by quantifying how large the day-to-night temperature oscillation is comparted to the summer-to-winter oscillation. Our results show a generally decreasing trend across the landscape (i.e., greater levels of annual temperature variability relative to an average day) over the past 40 years, while the variable’s importance as a driver increases. Precipitation seasonality is a measure of the variation in monthly precipitation totals over the course of the year. Our results show an increasing trend in precipitation seasonality as the variable itself remains a consistent model contributor. Taken together, the increasing and continuing importance of these two variables suggests that variability in both temperature levels and precipitation amounts has had a major historical influence on climatic suitability for Cassin’s Sparrow as well. The remaining, contributing variables both have a nexus with the monsoon season and drying conditions. In particular, precipitation of the wettest month (MC_Bio13) generally occurs during the monsoon season and the maximum temperature of the warmest month (MC-Bio05) generally precedes the onset of the monsoon season. Both of these variables show an increasing trend over the past 40 years, and both are consistent, low-level drivers in our results.

This interpretation of the importance of drying trends also appears to be supported by the patterns we see in the M2 time series, just at a different scale. In particular, the top contributing variables in the MC time series provide insights into the way macroclimatic change has potentially influenced seasonal trends while the most important variables in the M2 time series address various microclimatic and ecological functioning aspects of surface energy and hydrological dynamics (Table 6, Fig 6) [34,35,169,170]. The two most important variables in the M2 time series are surface wind speeds (SPEED), which appears to have been generally increasing across the south southwestern region of the study area over the past 40 years while remaining a consistently top model contributor, and bare soil evaporation energy flux (EVPSOIL), which is generally decreasing across the same area over this period while also remaining a top contributor across the time series. Wind has a mixing effect on the air near the ground surface that can increase the evaporation of water from soil and plant surfaces until a point is reached where drying conditions take over, as it appears to have occurred for the soil but not yet for plants. Water loss from plant surfaces, reflected as interception loss energy flux (EVPINTR), the fourth most contributory variable, has been increasing over the past 40 years while its importance as a driver in our models has steadily increased. A similar pattern is seen in the sixth most contributory variable, total re-evaporation of precipitation (PREVTOT), which is a measure of precipitation efficiency (i.e., the amount of falling precipitation that evaporates or sublimates before reaching the ground). In our results, there has been an overall increasing trend in PREVTOT across the study area while the driver itself has remained a moderately consistent in importance across the 40-year span of the study. These variables, along with increasing shortwave radiant energy from the sun (SWGNT), modulated by variably decreasing clouds across the North American Southwest (TAUTOT), contribute to observed conditions of absolute humidity (QV2M), which, on balance, at this point, have been generally increasing across the study area while decreasing in importance as an environmental driver [171–173]. Taken together, the top contributing variables in the M2 time series suggest that microclimatic drying, with more evaporation and less rainfall reaching the ground, may be the particular aspects of the varying monsoon precipitation pattern of importance to Cassin’s Sparrow.

This observed connection between top contributing variables, monsoon rainfall, and ground-level drying conditions is consistent with what is known about Cassin’s Sparrow’s ground-dwelling habit and the importance of microclimatic conditions to almost all aspects of the species’ life. Our results are also consistent with research showing that increasing temperatures, increased temperature and precipitation variability, and drying soils are potent drivers of environmental suitability at scales germane to many species found across the arid grasslands of the southwestern U.S. [174,175]. However, given the importance of Cassin’s Sparrow’s areal flight song behavior to the species’ breeding ecology, the increase in surface wind speed may have the most pronounced impact on climatic suitability and the species’ demography overall: male Cassin’s Sparrows simply do not skylark when it is windy [17].

What then are the take-away lessons from our analysis of environmental variables? For one, as noted by others, the juxtaposition of trend perspectives across two collections of predictors seems to provide a more holistic view of climate-species interactions than might be obtained from either collection on its own [49,55]. What is more, the automated, observation-guided, time-specific selection of variables appears to provide an integrated view across the two time series: the two collections of predictors seem to tell different parts of the same story about what might be driving change.

That being said, these results must be interpreted with caution. M2 variables represent the low-level physical drivers of many of the Earth system’s biological processes [27,176]: an interpretation of ecological plausibility could be made for many of the M2 variables. Studies have shown, however, that MaxEnt’s ranking of variable importance can capture biologically realistic assessments of factors governing range boundaries when models are built using best-practice procedures and variables are ranked based on permutation importance [69,79]. With Cassin’s Sparrow, we have a species whose behavioral and energetic ecology has been studied in detail, facilitating the interpretation of relationships with M2 variables [17]. Of the many potential contributors in the M2 and MC collections, the top contributing variables identified in the two time series are known to be important environmental influences for the species and are consistent with our mechanistic, process-based understanding of the bird’s natural history [2,17,177–179]. At this point, we feel confident saying that, in future modeling efforts, spatiotemporal attributes of North American Monsoon precipitation, hydrologic conditions of the soil, and surface winds should be considered variables of particular ecological relevance to Cassin’s Sparrow and key to understanding the species’ conservation status.

## Conclusions

This is a preliminary study, much of it qualitative, and all of it based on an experimental modeling protocol and suite of technologies that are still very much in development. While we see useful outcomes in what we have described here, our single biggest caution is that more work needs to be done to elevate overall confidence in the approach. We need experience applying retrospective ENM to different species, species assemblages, and different taxonomic groups. We also we need experience using larger and more diverse predictor collections as the source pool for variable screening, such as including a full range of topographic and edaphic data, vegetation indices, remotely sensed EFAs, and data on land-cover/land-use change.

Throughout this effort, we have tried to strike a balance between precision and scalability. It is likely that greater model accuracy across our time series could be attained by carefully considered variable selection and manual tuning. However, our workflow can be fully automated: the study presents a scalable approach to evaluating historical climatic suitability trends for species of conservation concern that has real potential to advance future species and habitat conservation activities.

For example, New Mexico, only one of the states occupied by Cassin’s Sparrow, is home to more than 5,800 species, about 235 of which have been identified as Species of Greatest Conservation Need [9,180,181]. In the current study, we looked at only one species. Even here, with only 30 and 19 variables respectively in the M2 and MC working set collections, at 100 samples per variable performed in triplicate, MERRA/Max selection across the two time series required 7,350 independent bivariate sampling runs and a total run time of about 90 minutes in our 100-core testbed. Run time on a single processor for this computational load would have been about four days. In theory, selection would require less than one minute in a fully provisioned 7,350-core cluster computing environment, which makes an otherwise intractable manual analysis possible and perhaps even scalable to accommodate comprehensive, state-level analysis of hundreds of species. It is not unreasonable to imagine a fully realized, operational implementation of this analytic workflow being made available to the research community on a cost-effective basis through one of many commercial cloud services [182–187].

There are methodological details in the current study that could raise concern, many of which we hope to address as our work in this area progresses. For example, we started with birds because of the ready access to historical, georeferenced occurrence and survey data on birds that are available through resources such as GBIF and USGS’s BBS. Application programing interfaces (APIs) and R libraries are making these resources increasingly easy to work with, but the vetting of observations, winnowing of records, and otherwise perfecting of ENM inputs remains a science based largely on expert knowledge, subject to controversy, and difficult to automate. Our default settings seem to work well for our purposes [74], but could be inadequate for other applications. It will be a challenge to extend this approach to organisms lacking the abundant, publicly-accessible observational records that are currently available for birds.

We are sometimes criticized for using reanalysis data for ENM studies, generally by numerical climate modelers who value reanalyses for different reasons and have reservations about using the data in other applications. Their concerns usually revolve around the potential biases or errors that might exist in what are primarily research datasets. However, we still see untapped value in these data because of their diversity of modeled attributes, the way reanalyses integrate such a wide range of Earth system measurements, and their remarkably high temporal resolution, which opens doors to a level of temporally fine-grained analysis difficult to obtain with other types of data. Furthermore, our focus is on the long-running patterns and trends in environmental drivers, which tend to accommodate bias and resolution issues that may pose problems in applications that rely on actual values from reanalysis datasets.

Attitudes appear to be evolving on this issue. For example, Copernicus, the Earth observation component of the European Union’s space program, now provides a suite of API-accessible, downscaled reanalysis and projection data products targeted for and vetted by the biodiversity research community [25,188–191]. Copernicus’s Climate Data Store [188] will make this type of work far more accessible, and the ability to anchor climatic suitability projections along a continuum that extends from historical trends to the future within a consistent information space could contribute an important new level of insight to conservation status assessments.

From a science perspective, the most significant limitation of the current work is that, at this point, it raises more questions than it answers. The ambiguities and uncertainties regarding the conservation status of Cassin’s Sparrow that inspired the project in the first place remain unresolved. However, these unanswered questions have helped us identify the key issues to be addressed in next steps. Four areas stand out as being particularly promising topics for future research. The first two directly address the ambiguities question; pursuing them will undoubtedly lead to useful insights into the natural history of Cassin’s Sparrow:

1. *State-level conservation status*. Given the variability we observed in climatic suitability across Cassin’s Sparrow’s range, along with the disparities noted between suitability- and abundance-derived perspectives of this species’ status, an updated, state-by-state look at the species’ distribution and conservation status is needed in order to better understand regional vulnerabilities and patterns of change. Such an exercise would also be an opportunity to examine the potential of deriving a useful climate change risk metric from retrospective ENM trends that could improve species vulnerability assessments across taxa [7,192–195].
2. *Interannual variability*. Given the variability we observed in climatic suitability across areas occupied by Cassin’s Sparrow during the breeding season, a closer look at the nature and scope of historical, interannual range changes is need. Combining M2 and MC predictors while separating breeding and non-breeding occurrences into monthly time series could potentially identify life-cycle-specific drivers influencing this variability and provide useful insights into the species’ itinerant breeding habit, recruitment, dispersal, and overall response to seasonal rainfall and surface wind speed patterns. It would also help address the open research questions raised by points (2) and (3) in the discussion. The next two topics are research questions that have grown out of the current work. While rooted in our interest in Cassin’s Sparrow, insights gained through exploration of these topics would likely be more generally applicable across species:
3. *Bridging the correlative-mechanistic (CM) divide*. Correlative ENM focuses on understanding the occurrence-based environmental associations that allow persistence of a species’ population; mechanistic modeling seeks to do the same, but based on first principles of biophysics and physiology [196]. The mechanistic links between climate and the environmental responses of an organism occur through the microclimatic conditions that organisms experience on the ground [51]. A growing body of work attests to the value of including biologically-relevant microclimatic variables alongside traditional, macroclimatic drivers in ENM [30,43,49,61,174,175], the practical upshot of which is improved results that blend the best of correlative and mechanistic insight [51,52,197–200]. A major challenge to doing so, however, has been the paucity of detailed knowledge about most organisms’ microclimatic and functional requirements, which are needed to parameterize mechanistic models [49,196–198,201]. That challenge notwithstanding, there appears to be little dispute about the types of drivers that represent microclimatic conditions of importance for a diversity of species: they include data on soil properties, wind speed, solar radiation, humidity, cloud cover, etc., essentially a subset of the M2 collection [51,196]. In the current study, we are struck by the apparent capacity of reanalysis data plus time-specific variable selection across a retrospective time series to simultaneously identify meaningful macroclimatic variables, microclimatic variables, and variables representing ecological function. This observation suggests the possibility that the approach we have taken here is a step toward bridging the CM divide. Given that Cassin’s Sparrow is one of a relatively small group of birds for which the mechanistic drivers behind many behavioral traits have been studied [17], the species could provide a useful focus for examining this issue further.
4. *Bridging the abundance-suitability (AS) divide*. We are intrigued by the apparent congruence between our centroid results and those of Huang et al. [124]: the velocity and direction of shifts in climatic suitability observed by Huang and his colleagues parallels the patterns we observed in our MC results; the velocity and direction of shifts in abundance observed by Huang parallels the results we observed in our M2 results. The similarities raise the question, in our minds, of whether reanalysis-based, retrospective environmental suitability analyses that combine both time-specificity and variable-specificity, as we have done in this study, might provide a practical means of estimating abundance distributions at scale using readily-available occurrence data. Work on this question would help address the open research question raised by point (3) of the discussion above as well as advance the management-relevant benefits suggested by point (4) of the discussion.

Finally, a pragmatic next step that will be important in order to address these research topics and enable the approach we have used here to become more generally applicable. We have essentially been exploring the capacity of automated, retrospective ENM to provide a new and useful dimension to studies of the temporal dynamics of climatic suitability. To gain experience and confidence with retrospective ENM, broader community engagement with these methods is needed. To that end, we have begun making the software and data supporting this research available as the MMX Toolkit [202]. We see continued development of the MMX Toolkit, accompanied by community contributions and refinement and feedback on work such as that described here, as key to maturing the methodology, making it truly useful to wildlife conservation activities, and increasing what we know about the current status of Cassin’s Sparrow and what the future might hold for this and other species.

## Appendix A. MERRA-2 (M2) variable definitions.*

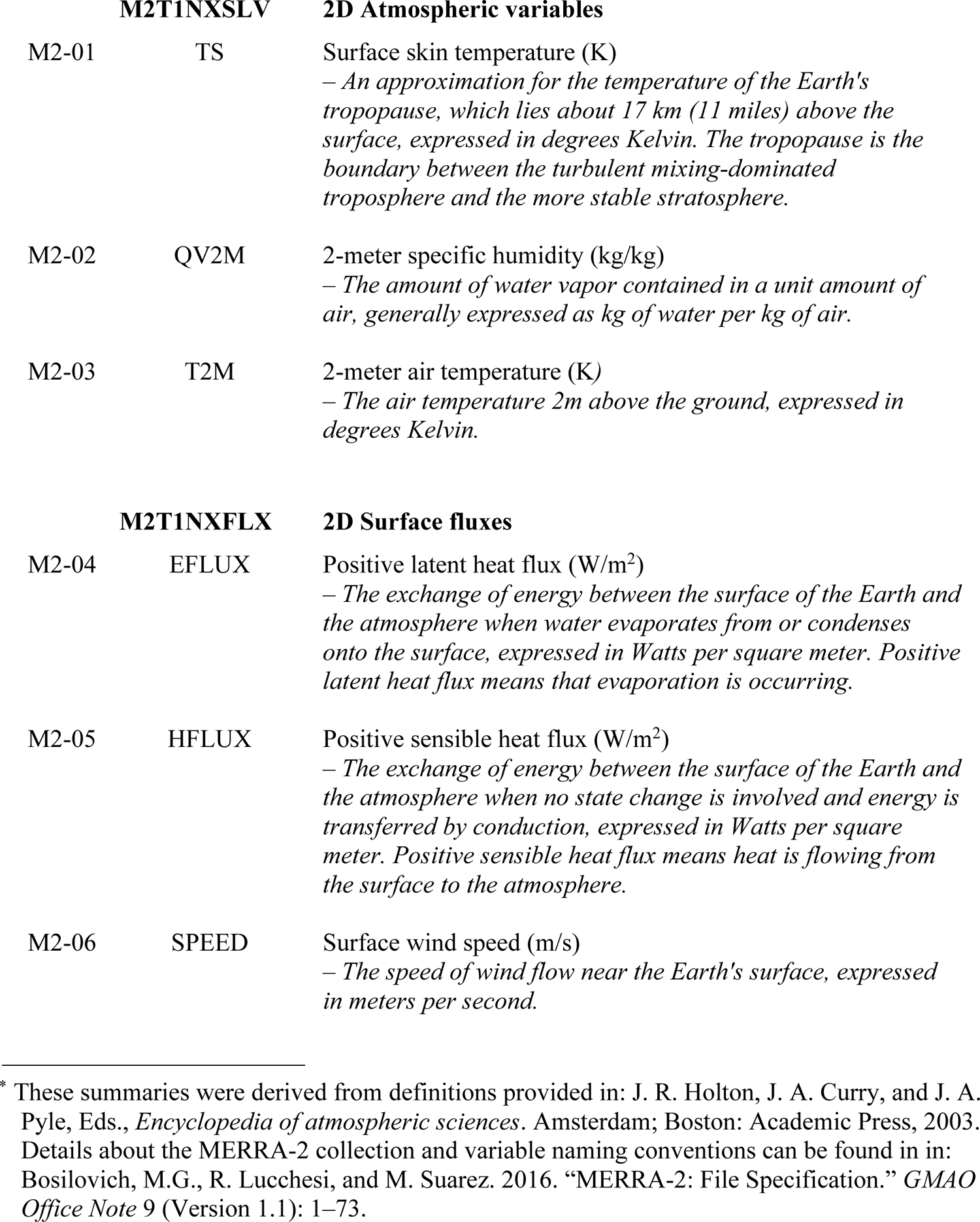
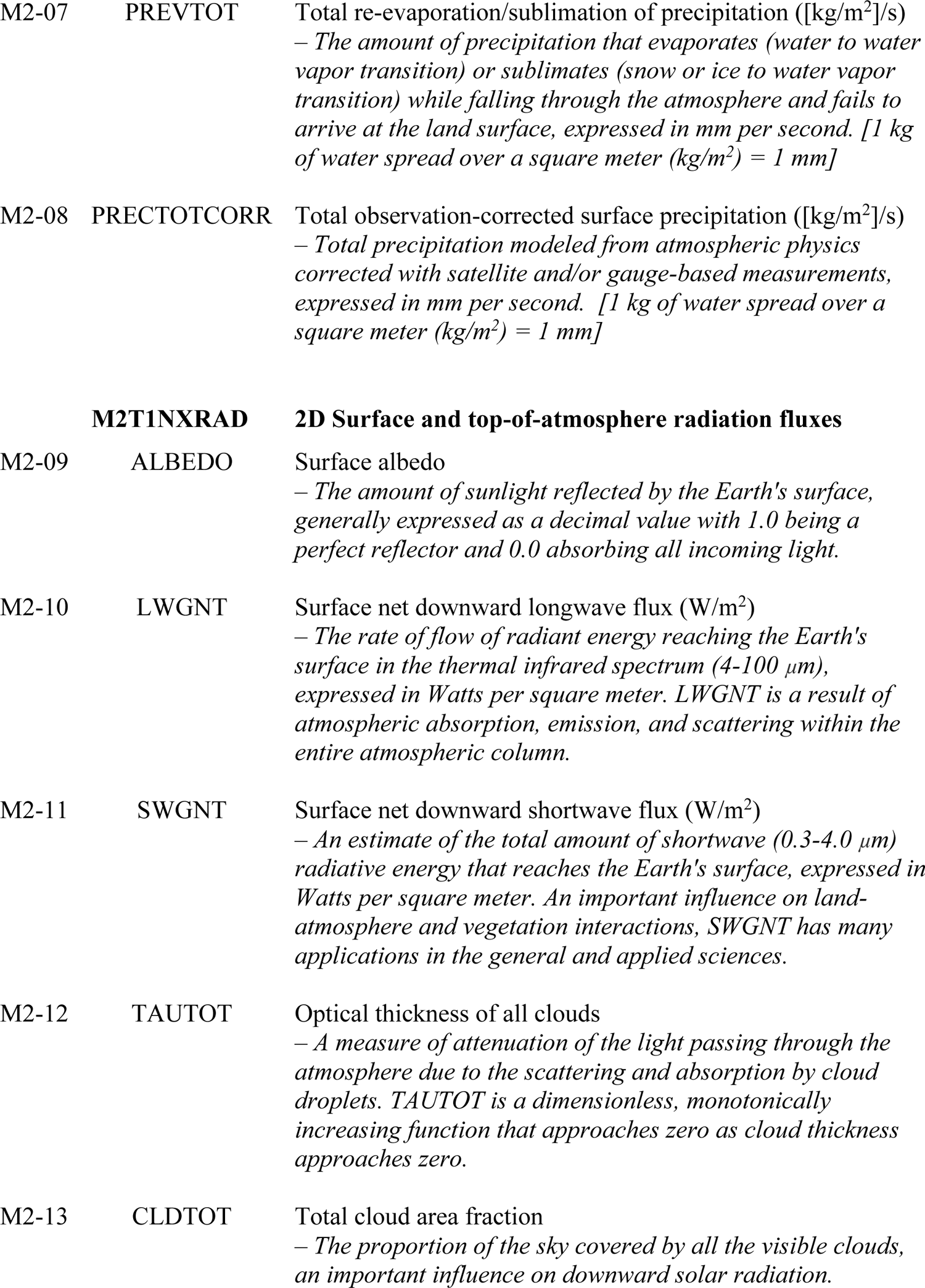
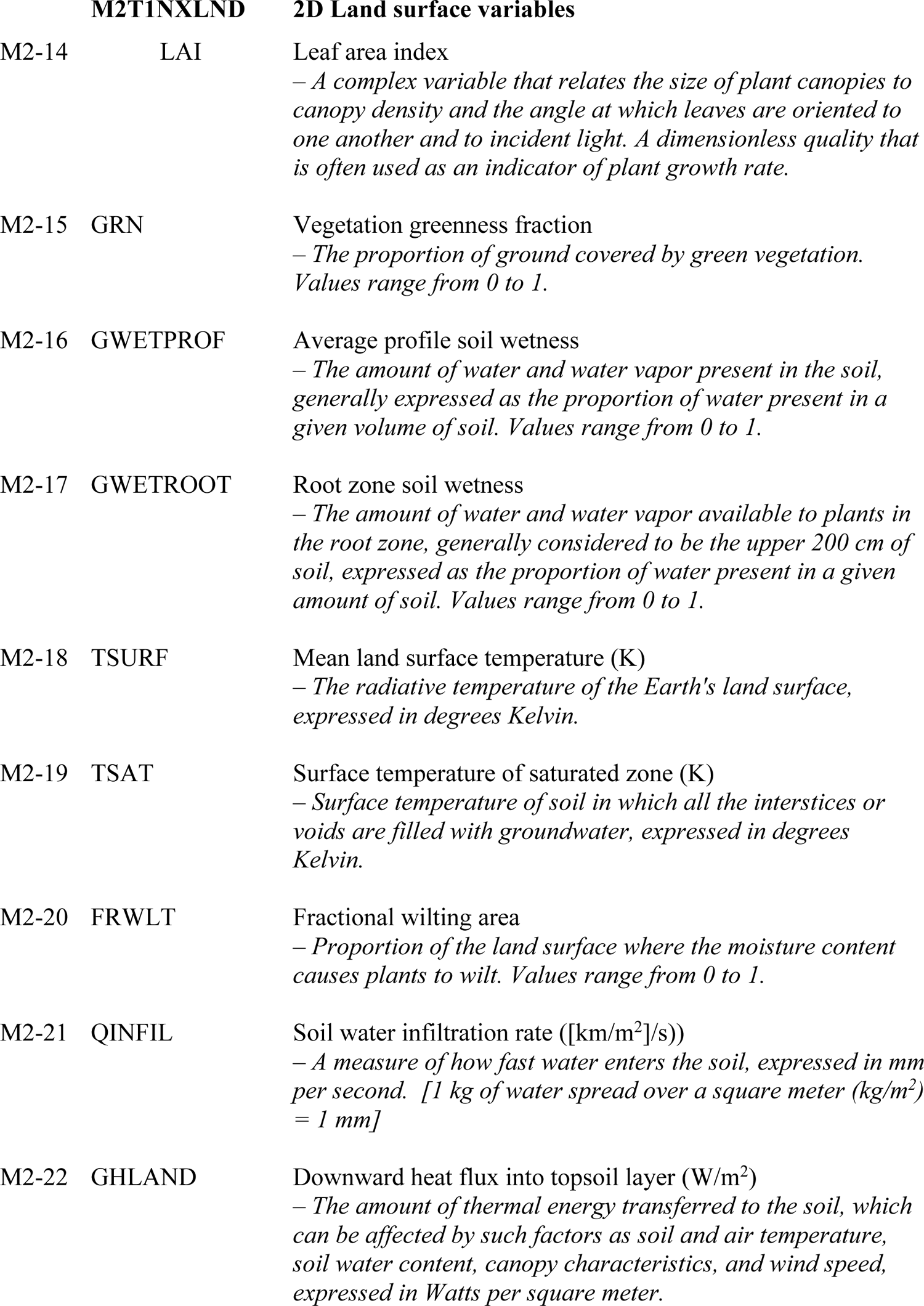
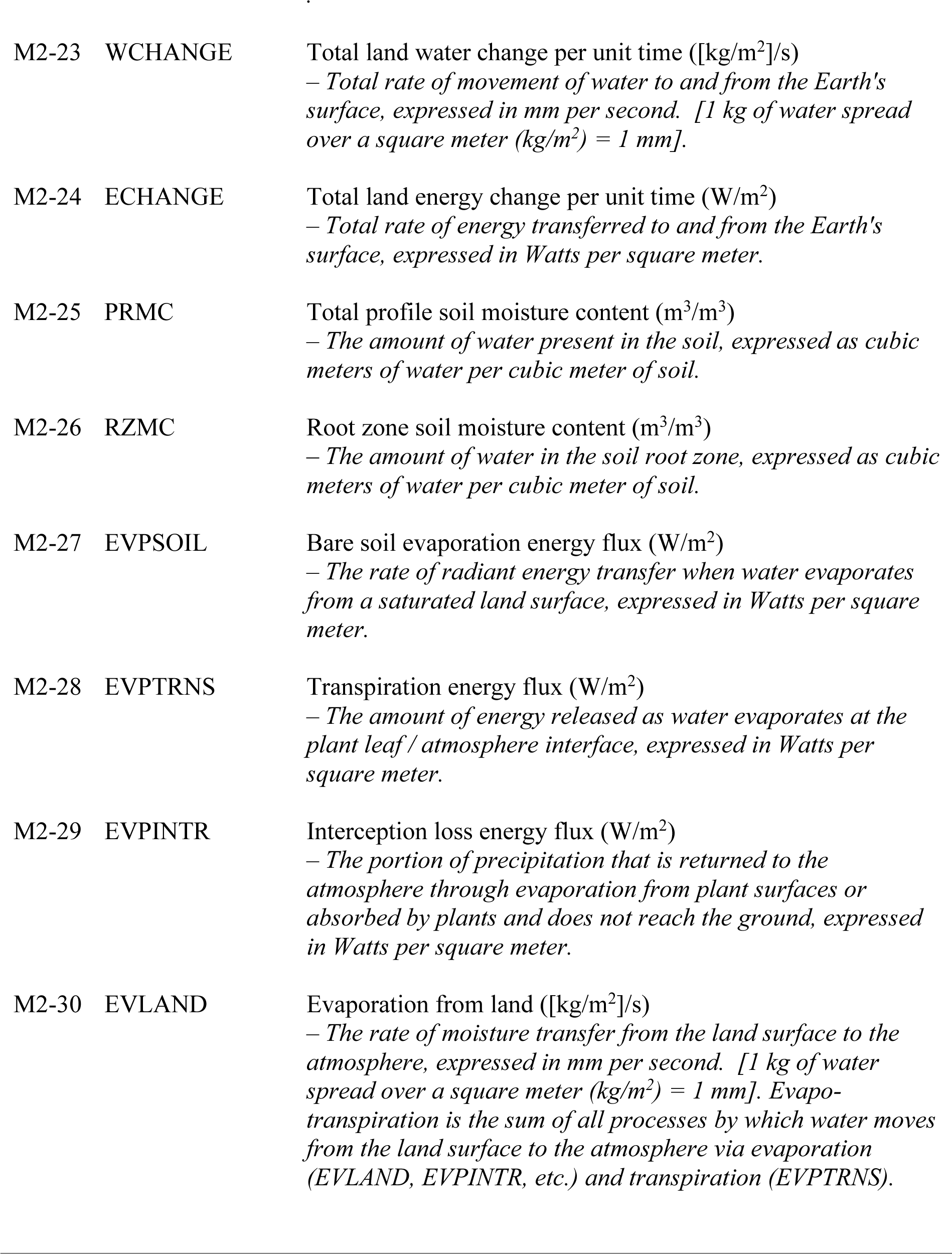

## Literature Cited

1. Woodhouse SW. Zonotrichia Cassinii, nobis. Proceedings of the Academy of Natural Science of Philadelphia. Philadelphia: Merrihow and Thompson; 1852. pp. 60–61. Available: https://www.biodiversitylibrary.org/item/17888#page/7/mode/1up

2. Dunning, Jr. JB, Bowers, Jr. RK, Suter SJ, Bock CE. Cassin’s Sparrow (Peucaea cassinii), Version 1.0. In: Birds of the World (P. G. Rodewald, Editor) [Internet]. 2020 [cited 23 Dec 2022]. Available: 10.2173/bow.casspa.01

3. Williams FC, LeSassier AL. Cassin’s Sparrow. In: Austin OL, editor. Life Histories of North American Cardinals, Grosbeaks, Buntings, Towhees, Finches, Sparrows, and Allies, Order Passeriformes, Family Fringillidae: (in 3vols) Part 2, Genera Pipilo (part) Through Spizella. New York: Dover; 1968. pp. 981–990.

4. Iknayan KJ, Beissinger SR. Collapse of a desert bird community over the past century driven by climate change. Proc Natl Acad Sci USA. 2018;115: 8597. doi:10.1073/pnas.1805123115

5. Ruth JM. Cassin’s Sparrow (Aimophila cassinii) status assessment and conservation plan. Denver, CO; 2000. Report No.: BTP-R6002-2000. Available: http://pubs.er.usgs.gov/publication/2002055

6. Wolf B. Global warming and avian occupancy of hot deserts; a physiological and behavioral perspective. Revista Chilena de Historia Natural. 2000;73: 395–400.

7. Wilsey C, Taylor L, Bateman B, Jensen C, Michel N, Panjabi A, et al. Climate policy action needed to reduce vulnerability of conservation-reliant grassland birds in North America. Conservat Sci and Prac. 2019;1. doi:10.1111/csp2.21

8. Bateman BL, Pidgeon AM, Radeloff VC, VanDerWal J, Thogmartin WE, Vavrus SJ, et al. The pace of past climate change vs. potential bird distributions and land use in the United States. Global Change Biology. 2016;22: 1130–1144. doi:10.1111/gcb.13154

9. Game NM, Fish. New Mexico State Wildlife Action Plan (SWAP) Final 2019. In: New Mexico Department of Game & Fish [Internet]. 2019 [cited 16 Oct 2022]. Available: https://www.wildlife.state.nm.us/wpfb-file/new-mexico-state-wildlife-action-plan-swap-final-2019-pdf/

10. Cassin’s Sparrow - Whatbird.com. [cited 22 May 2021]. Available: https://identify.whatbird.com/obj/278/overview/cassins_sparrow.aspx

11. Cassin’s Sparrow. In: Audubon [Internet]. 2014 [cited 22 May 2021]. Available: https://www.audubon.org/field-guide/bird/cassins-sparrow

12. Cassin’s Sparrow (Peucaea cassinii) - BirdLife species factsheet. [cited 22 May 2021]. Available: http://datazone.birdlife.org/species/factsheet/22721272

13. Cassin’s Sparrow Life History, All About Birds, Cornell Lab of Ornithology. [cited 22 May 2021]. Available: https://www.allaboutbirds.org/guide/Cassins_Sparrow/lifehistory

14. New Mexico Avian Conservation Partners | NMACP. [cited 7 Oct 2022]. Available: https://avianconservationpartners-nm.org/

15. Salas EAL, Seamster VA, Boykin KG, Harings NM, Dixon KW, Department of Fish, Wildlife and Conservation Ecology, New Mexico State University, Las Cruces, New Mexico 88003, USA. Modeling the impacts of climate change on Species of Concern (birds) in South Central U.S. based on bioclimatic variables. AIMS Environmental Science. 2017;4: 358–385. doi:10.3934/environsci.2017.2.358

16. Peucaea cassinii | NatureServe Explorer. [cited 16 Jun 2023]. Available: https://explorer.natureserve.org/Taxon/ELEMENT_GLOBAL.2.102862/Peucaea_cassinii

17. Schnase JL, Grant WE, Maxwell TC, Leggett JJ. Time and energy budgets of Cassin’s sparrow (Aimophila cassinii) during the breeding season: evaluation through modelling. Ecological Modelling. 1991;55: 285–319.

18. Lynn J. Cassin’s Sparrow (Aimophila cassinii): A Technical Conservation Assessment. USDA Forest Service; 2006. Available: http://www.fs.usda.gov/Internet/FSE_DOCUMENTS/stelprdb5182053.pdf

19. Maxwell TC. Avifauna of the Concho Valley of West-Central Texas with Special Reference to Historical Change. Texas A&M University. 1979.

20. Ohmart RD. Dual breeding ranges in Cassin’s sparrow (Aimophila cassinii). In: Hoff CC, Riedesel ML, editors. Physiological systems in semiarid environments. Albuquerque, NM: University of New Mexico Press; 1969. p. 105.

21. Schnase JL, Maxwell TC. Use of song patterns to identify individual male Cassin’s Sparrows. Journal of Field Ornithology. 1989;60: 12–19.

22. Maguire Jr B. Niche response structure and the analytical potentials of its relationship to the habitat. The American Naturalist. 1973;107: 213–246.

23. Pecl GT, Araújo MB, Bell JD, Blanchard J, Bonebrake TC, Chen I-C, et al. Biodiversity redistribution under climate change: Impacts on ecosystems and human well-being. Science. 2017;355: eaai9214. doi:10.1126/science.aai9214

24. Johnston A, Auer T, Fink D, Strimas-Mackey M, Iliff M, Rosenberg KV, et al. Comparing abundance distributions and range maps in spatial conservation planning for migratory species. Ecol Appl. 2020;30. doi:10.1002/eap.2058

25. Climate reanalysis | Copernicus. [cited 15 Oct 2022]. Available: https://climate.copernicus.eu/climate-reanalysis

26. Edwards PN. A Vast Machine: Computer Models, Climate Data, and the Politics of Global Warming. Cambridge, MA: MIT Press; 2010.

27. Cavanagh RD, Murphy EJ, Bracegirdle TJ, Turner J, Knowland CA, Corney SP, et al. A Synergistic Approach for Evaluating Climate Model Output for Ecological Applications. Frontiers in Marine Science. 2017;4: 308. doi:10.3389/fmars.2017.00308

28. Araújo MB, Anderson RP, Barbosa AM, Beale CM, Dormann CF, Early R, et al. Standards for distribution models in biodiversity assessments. Science Advances. 2019;5: eaat4858.

29. Barbet-Massin M, Jetz W. A 40-year, continent-wide, multispecies assessment of relevant climate predictors for species distribution modelling. Heikkinen R, editor. Diversity and Distributions. 2014;20: 1285–1295. doi:10.1111/ddi.12229

30. Leitão PJ, Santos MJ. Improving models of species ecological niches: A remote sensing overview. Front Ecol Evol. 2019;7: 9. doi:10.3389/fevo.2019.00009

31. Simões MVP, Peterson AT. Importance of biotic predictors in estimation of potential invasive areas: the example of the tortoise beetle *Eurypedus nigrosignatus*, in Hispaniola. PeerJ. 2018;6: e6052. doi:10.7717/peerj.6052

32. Petitpierre B, Broennimann O, Kueffer C, Daehler C, Guisan A. Selecting predictors to maximize the transferability of species distribution models: lessons from cross-continental plant invasions: Which predictors increase the transferability of SDMs? Global Ecology and Biogeography. 2017;26: 275–287. doi:10.1111/geb.12530

33. Austin MP, Van Niel KP. Improving species distribution models for climate change studies: variable selection and scale: Species distribution models for climate change studies. Journal of Biogeography. 2011;38: 1–8. doi:10.1111/j.1365-2699.2010.02416.x

34. Bosilovich MG, Lucchesi R, Suarez M. MERRA-2: File Specification. GMAO Office Note. 2016;9: 1–73.

35. Gelaro R, Mccarty W, Su MJ, Todling R, Molod A, Takacs L, et al. The Modern-Era Retrospective Analysis for Research and Applications, Version 2 (MERRA-2). JOURNAL OF CLIMATE. 2017;30: 36.

36. Fick SE, Hijmans RJ. WorldClim 2: new 1-km spatial resolution climate surfaces for global land areas. International Journal of Climatology. 2017;37: 4302–4315. doi:10.1002/joc.5086

37. Worldclim bioclimatic variables. 2020 [cited 22 May 2020]. Available: https://worldclim.org/data/worldclim21.html

38. Beaumont LJ, Hughes L, Poulsen M. Predicting species distributions: use of climatic parameters in BIOCLIM and its impact on predictions of species’ current and future distributions. Ecological Modelling. 2005;186: 251–270. doi:10.1016/j.ecolmodel.2005.01.030

39. Booth TH, Nix HA, Busby JR, Hutchinson MF. BIOCLIM: the first species distribution modelling package, its early applications and relevance to most current MaxEnt studies. Franklin J, editor. Diversity Distrib. 2014;20: 1–9. doi:10.1111/ddi.12144

40. Hijmans RJ, Cameron SE, Parra JL, Jones PG, Jarvis A. Very high resolution interpolated climate surfaces for global land areas. International Journal of Climatology. 2005;25: 1965–1978. doi:10.1002/joc.1276

41. O’Donnell MS, Ignizio D a. Bioclimatic Predictors for Supporting Ecological Applications in the Conterminous United States. Reston, VA: US Geological Survey; 2012 p. 10. Report No.: 691. Available: https://pubs.usgs.gov/ds/691/

42. Vega GC, Pertierra LR, Olalla-Tárraga MÁ. Data from: MERRAclim, a high-resolution global dataset of remotely sensed bioclimatic variables for ecological modelling. Dryad; 2018. p. 8324359732 bytes. doi:10.5061/DRYAD.S2V81

43. Cabello J, Fernández N, Alcaraz-Segura D, Oyonarte C, Piñeiro G, Altesor A, et al. The ecosystem functioning dimension in conservation: insights from remote sensing. Biodivers Conserv. 2012;21: 3287–3305. doi:10.1007/s10531-012-0370-7

44. Porzig EL, Seavy NE, Gardali T, Geupel GR, Holyoak M, Eadie JM. Habitat suitability through time: using time series and habitat models to understand changes in bird density. Ecosphere. 2014;5: art12. doi:10.1890/ES13-00166.1

45. Gonçalves J, Alves P, Pôças I, Marcos B, Sousa-Silva R, Lomba Â, et al. Exploring the spatiotemporal dynamics of habitat suitability to improve conservation management of a vulnerable plant species. Biodivers Conserv. 2016;25: 2867–2888. doi:10.1007/s10531-016-1206-7

46. Brooks TM, Pimm SL, Akçakaya HR, Buchanan GM, Butchart SHM, Foden W, et al. Measuring terrestrial area of habitat (AOH) and its utility for the IUCN Red List. Trends in Ecology & Evolution. 2019;34: 977–986. doi:10.1016/j.tree.2019.06.009

47. Oeser J, Heurich M, Senf C, Pflugmacher D, Belotti E, Kuemmerle T. Habitat metrics based on multi-temporal Landsat imagery for mapping large mammal habitat. Pettorelli N, Armenteras D, editors. Remote Sens Ecol Conserv. 2020;6: 52–69. doi:10.1002/rse2.122

48. Regos A, Gómez-Rodríguez P, Arenas-Castro S, Tapia L, Vidal M, Domínguez J. Model-assisted bird monitoring based on remotely sensed ecosystem functioning and atlas data. Remote Sensing. 2020;12: 2549. doi:10.3390/rs12162549

49. Regos A, Gonçalves J, Arenas-Castro S, Alcaraz-Segura D, Guisan A, Honrado JP. Mainstreaming remotely sensed ecosystem functioning in ecological niche models. He K, Moat J, editors. Remote Sens Ecol Conserv. 2022;8: 431–447. doi:10.1002/rse2.255

50. Arenas-Castro S, Sillero N. Cross-scale monitoring of habitat suitability changes using satellite time series and ecological niche models. Science of The Total Environment. 2021;784: 147172. doi:10.1016/j.scitotenv.2021.147172

51. Kearney MR, Isaac AP, Porter WP. microclim: Global estimates of hourly microclimate based on long-term monthly climate averages. Sci Data. 2014;1: 140006. doi:10.1038/sdata.2014.6

52. Kearney M, Simpson SJ, Raubenheimer D, Helmuth B. Modelling the ecological niche from functional traits. Phil Trans R Soc B. 2010;365: 3469–3483. doi:10.1098/rstb.2010.0034

53. Lembrechts JJ, Nijs I, Lenoir J. Incorporating microclimate into species distribution models. Ecography. 2019;42: 1267–1279. doi:10.1111/ecog.03947

54. Guisan A, Tingley R, Baumgartner JB, Naujokaitis-Lewis I, Sutcliffe PR, Tulloch AIT, et al. Predicting species distributions for conservation decisions. Arita H, editor. Ecol Lett. 2013;16: 1424–1435. doi:10.1111/ele.12189

55. Ingenloff K, Peterson AT. Incorporating time into the traditional correlational distributional modelling framework: A proof-of-concept using the Wood Thrush *Hylocichla mustelina*. Peres-Neto P, editor. Methods Ecol Evol. 2021;12: 311–321. doi:10.1111/2041-210X.13523

56. Rollinson CR, Finley AO, Alexander MR, Banerjee S, Dixon Hamil K-A, Koenig LE, et al. Working across space and time: nonstationarity in ecological research and application. Front Ecol Environ. 2021;19: 66–72. doi:10.1002/fee.2298

57. Bede-Fazekas Á, Somodi I. The way bioclimatic variables are calculated has impact on potential distribution models. Freckleton R, editor. Methods Ecol Evol. 2020;11: 1559– 1570. doi:10.1111/2041-210X.13488

58. Ingenloff K, Hensz CM, Anamza T, Barve V, Campbell LP, Cooper JC, et al. Predictable invasion dynamics in North American populations of the Eurasian collared dove *Streptopelia decaocto*. Proc R Soc B. 2017;284: 20171157. doi:10.1098/rspb.2017.1157

59. Peterson AT, Martínez-Campos C, Nakazawa Y, Martínez-Meyer E. Time-specific ecological niche modeling predicts spatial dynamics of vector insects and human dengue cases. Transactions of The Royal Society of Tropical Medicine and Hygiene. 2005;99: 647–655. doi:10.1016/j.trstmh.2005.02.004

60. Ingenloff K. Enhancing the correlative ecological niche modeling framework to incorporate the temporal dimension of species’ distributions. University of Kansas. 2020.

61. Barve N, Martin C, Brunsell NA, Peterson AT. The role of physiological optima in shaping the geographic distribution of Spanish moss: Physiological optima of Spanish moss. Global Ecology and Biogeography. 2014;23: 633–645. doi:10.1111/geb.12150

62. Roubicek AJ, VanDerWal J, Beaumont LJ, Pitman AJ, Wilson P, Hughes L. Does the choice of climate baseline matter in ecological niche modelling? Ecological Modelling. 2010;221: 2280–2286. doi:10.1016/j.ecolmodel.2010.06.021

63. Harris RMB, Grose MR, Lee G, Bindoff NL, Porfirio LL, Fox-Hughes P. Climate projections for ecologists. Wiley Interdisciplinary Reviews: Climate Change. 2014;5: 621– 637. doi:10.1002/wcc.291

64. Goberville E, Beaugrand G, Hautekèete N-C, Piquot Y, Luczak C. Uncertainties in the projection of species distributions related to general circulation models. Ecol Evol. 2015;5: 1100–1116. doi:10.1002/ece3.1411

65. Pérez-Navarro MA, Broennimann O, Esteve MA, Moya-Perez JM, Carreño MF, Guisan A, et al. Temporal variability is key to modelling the climatic niche. Diversity and Distributions. 2021;27: 473–484. doi:10.1111/ddi.13207

66. Guida RJ, Abella SR, Jr WJS, Stephen H, Roberts CL. Climatic change and desert vegetation distribution: Assessing thirty years of change in southern nevada’s Mojave Desert. The Professional Geographer. 2014;66: 311–322. doi:10.1080/00330124.2013.787007

67. Guida RJ, Abella SR, Roberts CL, Norman CM, Smith Jr. WJ. Assessing historical and future habitat models for four conservation-priority Mojave Desert species. Journal of Biogeography. 2019;46: 2081–2097. 10.1111/jbi.13645

68. Araújo MB, Guisan A. Five (or so) challenges for species distribution modelling. Journal of Biogeography. 2006;33: 1677–1688. doi:10.1111/j.1365-2699.2006.01584.x

69. Smith AB, Santos MJ. Testing the ability of species distribution models to infer variable importance. Ecography. 2020;43: 1801–1813. doi:10.1111/ecog.05317

70. Cobos ME, Peterson AT, Osorio-Olvera L, Jiménez-García D. An exhaustive analysis of heuristic methods for variable selection in ecological niche modeling and species distribution modeling. Ecological Informatics. 2019;53: 100983. doi:10.1016/j.ecoinf.2019.100983

71. Peterson AT, Cobos ME, Jiménez-García D. Major challenges for correlational ecological niche model projections to future climate conditions: Climate change, ecological niche models, and uncertainty. Annals of the New York Academy of Sciences. 2018;1429: 66–77. doi:10.1111/nyas.13873

72. Peterson AT, Nakazawa Y. Environmental data sets matter in ecological niche modelling: an example with Solenopsis invicta and Solenopsis richteri. Global Ecology and Biogeography. 2008;0: 071113201427001-??? doi:10.1111/j.1466-8238.2007.00347.x

73. Zeng Y, Low BW, Yeo DCJ. Novel methods to select environmental variables in MaxEnt: A case study using invasive crayfish. Ecological Modelling. 2016;341: 5–13. 10.1016/j.ecolmodel.2016.09.019

74. Schnase JL, Carroll ML. Automatic variable selection in ecological niche modeling: A case study using Cassin’s Sparrow (Peucaea cassinii). PLOS ONE. 2022;17: e0257502. doi:10.1371/journal.pone.0257502

75. Schnase JL, Carroll ML, Gill RL, Tamkin GS, Li J, Strong SL, et al. Toward a Monte Carlo approach to selecting climate variables in MaxEnt. PLOS ONE. 2021;16: e0237208. doi:10.1371/journal.pone.0237208

76. Elith J, Phillips SJ, Hastie T, Dudík M, Chee YE, Yates CJ. A statistical explanation of MaxEnt for ecologists. Diversity and distributions. 2011;17: 43–57.

77. Phillips SJ, Anderson RP, Dudík M, Schapire RE, Blair ME. Opening the black box: An open-source release of Maxent. Ecography. 2017;40: 887–893.

78. Phillips SJ. A Brief Tutorial on Maxent. AT&T Research. 2005;190: 231–259.

79. Searcy CA, Shaffer HB. Do ecological niche models accurately identify climatic determinants of species ranges? The American Naturalist. 2016;187: 423–435. doi:10.1086/685387

80. Schnase JL, Carroll ML. The MMX Toolkit: High performance, reanalysis-based climatic suitability modeling to advance avian conservation. In: Albani S, Lumnitz S, Soille P, editors. Proceedings of the 2023 conference on Big Data from Space – 6-9 November 2023. Vienna: European Commission, Joint Research Centre, Publications Office; 2023. (to appear)

81. Chamberlain S, Oldoni D, Barve V, Desmet P, Geffert L, Mcglinn D, et al. rgbif: Interface to the Global Biodiversity Information Facility API. 2022. Available: https://CRAN.R-project.org/package=rgbif

82. GBIF - Global Biodiversity Information Facility. 2023 [cited 14 Aug 2023]. Available: https://www.gbif.org/

83. Cassin’s Sparrow - eBird. [cited 5 Feb 2022]. Available: https://ebird.org/species/casspa

84. iNaturalist. In: iNaturalist [Internet]. [cited 6 Oct 2022]. Available: https://www.inaturalist.org/

85. Data and Samples | NSF NEON | Open Data to Understand our Ecosystems. [cited 26 Dec 2022]. Available: https://www.neonscience.org/data-samples

86. Smithsonian National Museum of Natural History. [cited 26 Dec 2022]. Available: https://naturalhistory.si.edu/

87. Department of Ornithology | AMNH. In: American Museum of Natural History [Internet]. [cited 26 Dec 2022]. Available: https://bigbangblocks.amnh.org/research/vertebrate-zoology/ornithology

88. Museum of Comparative Zoology. [cited 26 Dec 2022]. Available: https://mcz.harvard.edu/home

89. Snow FH. Catalogue of the Birds of Kansas. Transactions of the Kansas Academy of Science (1872-1880). 1872;1: 21–29. doi:10.2307/3623510

90. University of Arizona Museum of Natural History. [cited 26 Dec 2022]. Available: http://eebweb.arizona.edu/Collections/

91. Boria RA, Blois JL. The effect of large sample sizes on ecological niche models: Analysis using a North American rodent, Peromyscus maniculatus. Ecological Modelling. 2018;386: 83–88. 10.1016/j.ecolmodel.2018.08.013

92. Boria RA, Olson LE, Goodman SM, Anderson RP. Spatial filtering to reduce sampling bias can improve the performance of ecological niche models. Ecological Modelling. 2014;275: 73–77. doi:10.1016/j.ecolmodel.2013.12.012

93. Sillero N, Barbosa AM. Common mistakes in ecological niche models. International Journal of Geographical Information Science. 2021;35: 213–226. doi:10.1080/13658816.2020.1798968

94. Kramer-Schadt S, Niedballa J, Pilgrim JD, Schröder B, Lindenborn J, Reinfelder V, et al. The importance of correcting for sampling bias in MaxEnt species distribution models. Robertson M, editor. Diversity Distrib. 2013;19: 1366–1379. doi:10.1111/ddi.12096

95. Gap Analysis Project | U.S. Geological Survey. [cited 10 Oct 2022]. Available: https://www.usgs.gov/programs/gap-analysis-project

96. Reichle RH. The MERRRA-Land Data Product. GMAO Office Note No 3 (Version 12). 2012; 38.

97. NASA Center for Climate Simulation | High Performance Computing for Science. [cited 6 Oct 2022]. Available: https://www.nccs.nasa.gov/

98. GES DISC - Goddard Earth Science Data and Information Services Center. [cited 26 May 2021]. Available: https://disc.gsfc.nasa.gov/

99. GES DISC Tools: MERRA Subsetter. [cited 6 Oct 2022]. Available: https://disc.gsfc.nasa.gov/information/tools?title=MERRA%20Subsetter

100. Hoyer S, Hamman JJ. xarray: N-D labeled Arrays and Datasets in Python. Journal of Open Research Software. 2017;5: 10. doi:10.5334/jors.148

101. GDAL/OGR Geospatial Data Abstraction Software Library. Open Source Geospatial Foundation; 2020. Available: https://gdal.org/

102. Hijmans RJ, Phillips S, Elith J, Leathwick J. dismo: Species Distribution Modeling. 2017. Available: https://CRAN.R-project.org/package=dismo

103. Hijmans RJ, van Etten J, Sumner M, Cheng J, Baston D, Bevan A, et al. raster: Geographic Data Analysis and Modeling. 2022. Available: https://CRAN.R-project.org/package=raster

104. Hijmans RJ, Phillips S, Elith JL and J. dismo: Species Distribution Modeling. 2022. Available: https://CRAN.R-project.org/package=dismo

105. C. Vega G, Pertierra LR, Olalla-Tárraga MÁ. MERRAclim, a high-resolution global dataset of remotely sensed bioclimatic variables for ecological modelling. Scientific Data. 2017;4. doi:10.1038/sdata.2017.78

106. Reichle RH, Liu Q, Koster RD, Draper CS, Mahanama SPP, Partyka GS. Land surface precipitation in MERRA-2. Journal of Climate. 2017;30: 1643–1664. doi:10.1175/JCLI-D-16-0570.1

107. Explore | NASA Center for Climate Simulation. [cited 17 Oct 2022]. Available: https://www.nccs.nasa.gov/systems/ADAPT

108. Pradhan P. Strengthening MaxEnt modelling through screening of redundant explanatory bioclimatic variables with variance inflation factor analysis. Researcher. 2016;8: 29–34.

109. Naimi B. usdm: Uncertainty Analysis for Species Distribution Models. 2017. Available: https://CRAN.R-project.org/package=usdm

110. Muscarella R, Galante PJ, Soley-Guardia M, Boria RA, Kass JM, Uriarte M, et al. ENMeval: An R package for conducting spatially independent evaluations and estimating optimal model complexity for Maxent ecological niche models. Methods in Ecology and Evolution. 2014;5: 1198–1205. doi:10.1111/2041-210X.12261

111. Kass JM, Muscarella R, Galante PJ, Bohl C, Buitrago-Pinilla GE, Boria RA, et al. ENMeval: Automated Tuning and Evaluations of Ecological Niche Models. 2022. Available: https://CRAN.R-project.org/package=ENMeval

112. mrmaxent. Maxent. 2022. Available: https://github.com/mrmaxent/Maxent

113. Phillips S. A Brief Tutorial on Maxent. 2010; 29.

114. H. Akaike. A new look at the statistical model identification. IEEE Transactions on Automatic Control. 1974;19: 716–723. doi:10.1109/TAC.1974.1100705

115. Phillips SJ, Anderson RP, Schapire RE. Maximum entropy modeling of species geographic distributions. Ecological Modelling. 2006;190: 231–259. doi:10.1016/j.ecolmodel.2005.03.026.

116. Phillips SJ, Research T. A Brief Tutorial on Maxent. 2017; 39.

117. Stephens PA, Mason LR, Green RE, Gregory RD, Sauer JR, Alison J, et al. Consistent response of bird populations to climate change on two continents. Science. 2016;352: 84–87. doi:10.1126/science.aac4858

118. Sen PK. Estimates of the regression coefficient based on Kendall’s tau. Journal of the American statistical association. 1968;63: 1379–1389.

119. Theil H. A rank-invariant method of linear and polynomial regression analysis. Indagationes mathematicae. 1950;12: 173.

120. Siegel AF. Robust regression using repeated medians. Biometrika. 1982;69: 242–244.

121. Evans JS, Murphy MA, Ram K. spatialEco: Spatial Analysis and Modelling Utilities. 2022. Available: https://CRAN.R-project.org/package=spatialEco

122. VanDerWal J, Murphy HT, Kutt AS, Perkins GC, Bateman BL, Perry JJ, et al. Focus on poleward shifts in species’ distribution underestimates the fingerprint of climate change. Nature Clim Change. 2013;3: 239–243. doi:10.1038/nclimate1688

123. Huang Q, Sauer JR, Swatantran A, Dubayah R. A centroid model of species distribution with applications to the Carolina wren *Thryothorus ludovicianus* and house finch *Haemorhous mexicanus* in the United States. Ecography. 2016;39: 54–66. doi:10.1111/ecog.01447

124. Huang Q, Bateman BL, Michel NL, Pidgeon AM, Radeloff VC, Heglund P, et al. Modeled distribution shifts of North American birds over four decades based on suitable climate alone do not predict observed shifts. Science of The Total Environment. 2023;857: 159603. doi:10.1016/j.scitotenv.2022.159603

125. La Sorte FA, Jetz W. Tracking of climatic niche boundaries under recent climate change: Niche tracking under recent climate change. Journal of Animal Ecology. 2012;81: 914– 925. doi:10.1111/j.1365-2656.2012.01958.x

126. Smith AB. enmSdmX: Species Distribution Modeling and Ecological Niche Modeling. 2023. Available: https://cran.r-project.org/web/packages/enmSdmX/index.html

127. Phillips SJ, Dudík M. Modeling of species distributions with Maxent: new extensions and a comprehensive evaluation. Ecography. 2008;31: 161–175.

128. (Peucaea cassinii) - Species Map - eBird. 2022 [cited 30 Sep 2022]. Available: https://ebird.org/map/casspa

129. Fielding AH, Bell JF. A review of methods for the assessment of prediction errors in conservation presence/absence models. Environmental Conservation. 1997;24: 38–49. doi:10.1017/S0376892997000088

130. Lobo JM, Jiménez-Valverde A, Real R. AUC: a misleading measure of the performance of predictive distribution models. Global Ecology and Biogeography. 2007; 7.

131. Allouche O, Tsoar A, Kadmon R. Assessing the accuracy of species distribution models: prevalence, kappa and the true skill statistic (TSS): Assessing the accuracy of distribution models. Journal of Applied Ecology. 2006;43: 1223–1232. doi:10.1111/j.1365-2664.2006.01214.x

132. Frans VF. True Skill Statistic (TSS) calculation across multiple maxent runs. Michigan State University; 2018 p. 5.

133. Warren DL, Seifert SN. Ecological niche modeling in Maxent: The importance of model complexity and the performance of model selection criteria. Ecological Applications. 2011;21: 335–342. doi:10.1890/10-1171.1

134. Arenas-Castro S, Gonçalves J, Alves P, Alcaraz-Segura D, Honrado JP. Assessing the multi-scale predictive ability of ecosystem functional attributes for species distribution modelling. Joseph S, editor. PLoS ONE. 2018;13: e0199292. doi:10.1371/journal.pone.0199292

135. Saunders SP, Michel NL, Bateman BL, Wilsey CB, Dale K, LeBaron GS, et al. Community science validates climate suitability projections from ecological niche modeling. Ecol Appl. 2020;30. doi:10.1002/eap.2128

136. Langham GM, Schuetz JG, Distler T, Soykan CU, Wilsey C. Conservation Status of North American Birds in the Face of Future Climate Change. LaDeau SL, editor. PLoS ONE. 2015;10: e0135350. doi:10.1371/journal.pone.0135350

137. Lehikoinen A, Virkkala R. North by north-west: climate change and directions of density shifts in birds. Glob Change Biol. 2016;22: 1121–1129. doi:10.1111/gcb.13150

138. North American Breeding Bird Survey. 2017. Available: https://www.pwrc.usgs.gov/bbs/.

139. Hubbard JP. The status of Cassin’s Sparrow in New Mexico and adjacent states. American Birds. 1977;31: 933–941.

140. Gillings S, Balmer DE, Fuller RJ. Directionality of recent bird distribution shifts and climate change in Great Britain. Global Change Biology. 2015;21: 2155–2168. 10.1111/gcb.12823

141. Bateman BL, Pidgeon AM, Radeloff VC, Flather CH, VanDerWal J, Akçakaya HR, et al. Potential breeding distributions of U.S. birds predicted with both short-term variability and long-term average climate data. Ecological Applications. 2016;26: 2720–2731. doi:10.1002/eap.1416

142. NatureServe Conservation Status Assessment | NatureServe. [cited 17 Jun 2023]. Available: https://www.natureserve.org/conservation-status-assessment

143. BBS - USGS Patuxent Wildlife Research Center. [cited 18 Jun 2023]. Available: https://www.pwrc.usgs.gov/bbs/

144. USFWS Association of Fish and Wildlife Agencies State Wildlife Action Plans (SWAP): Home. [cited 18 Jun 2023]. Available: https://www.fishwildlife.org/afwa-informs/state-wildlife-action-plans

145. Weber MM, Stevens RD, Diniz-Filho JAF, Grelle CEV. Is there a correlation between abundance and environmental suitability derived from ecological niche modelling? A meta-analysis. Ecography. 2017;40: 817–828. doi:10.1111/ecog.02125

146. Dallas TA, Hastings A. Habitat suitability estimated by niche models is largely unrelated to species abundance. Global Ecol Biogeogr. 2018;27: 1448–1456. doi:10.1111/geb.12820

147. Schofield LN, Siegel RB, Loffland HL. Modeling climate-driven range shifts in populations of two bird species limited by habitat independent of climate. Ecosphere. 2023;14. doi:10.1002/ecs2.4408

148. Cavalcante T, Weber MM, Barnett AA. Combining geospatial abundance and ecological niche models to identify high-priority areas for conservation: The neglected role of broadscale interspecific competition. Front Ecol Evol. 2022;10: 915325. doi:10.3389/fevo.2022.915325

149. Arroyo B, Estrada A, Casas F, Cardador L, De Cáceres M, Bota G, et al. Functional habitat suitability and urban encroachment explain temporal and spatial variations in abundance of a declining farmland bird, the Little Bustard Tetrax tetrax. ACE. 2022;17: art19. doi:10.5751/ACE-02243-170219

150. Monnier-Corbel A, Robert A, Hingrat Y, Benito BM, Monnet A-C. Species Distribution Models predict abundance and its temporal variation in a steppe bird population. Global Ecology and Conservation. 2023;43: e02442. doi:10.1016/j.gecco.2023.e02442

151. Meehan TD, Michel NL, Rue H. Spatial modeling of Audubon Christmas Bird Counts reveals fine-scale patterns and drivers of relative abundance trends. Ecosphere. 2019;10: e02707. doi:10.1002/ecs2.2707

152. Osorio-Olvera LA, Falconi M, Soberón J. Sobre la relación entre idoneidad del hábitat y la abundancia poblacional bajo diferentes escenarios de dispersión. Revista Mexicana de Biodiversidad. 2016;87: 1080–1088. doi:10.1016/j.rmb.2016.07.001

153. Lunghi E, Manenti R, Mulargia M, Veith M, Corti C, Ficetola GF. Environmental suitability models predict population density, performance and body condition for microendemic salamanders. Sci Rep. 2018;8: 7527. doi:10.1038/s41598-018-25704-1

154. Alcaraz-Segura D, Lomba A, Sousa-Silva R, Nieto-Lugilde D, Alves P, Georges D, et al. Potential of satellite-derived ecosystem functional attributes to anticipate species range shifts. International Journal of Applied Earth Observation and Geoinformation. 2017;57: 86–92. doi:10.1016/j.jag.2016.12.009

155. Bradie J, Leung B. A quantitative synthesis of the importance of variables used in MaxEnt species distribution models. Journal of Biogeography. 2017;44: 1344–1361. doi:10.1111/jbi.12894

156. Currie DJ, Venne S. Climate change is not a major driver of shifts in the geographical distributions of North American birds. Global Ecology and Biogeography. 2017;26: 333– 346.

157. Harris DB, Gregory SD, Brook BW, Ritchie EG, Croft DB, Coulson G, et al. The influence of non-climate predictors at local and landscape resolutions depends on the autecology of the species: Autecology informs SDM scale and predictor choice. Austral Ecology. 2014;39: 710–721. doi:10.1111/aec.12134

158. Schnase JL, Carroll ML. MERRA/Max: Harnessing the Potential of Climate Model Outputs in Studies of Ecosystem Change. ESA 2020 Virtual Meeting. ESA; 2020. Available: https://eco.confex.com/eco/2020/meetingapp.cgi/Paper/86032

159. Carroll ML, Schnase JL, Gill RL, Tamkin GS, Li J, Maxwell TP, et al. MERRAMax: A machine learning approach to stochastic convergence with a multi-variate dataset. IGARSS 2020 - 2020 IEEE International Geoscience and Remote Sensing Symposium. Waikoloa, HI, USA: IEEE; 2020. pp. 2017–2020. doi:10.1109/IGARSS39084.2020.9324185

160. Carroll ML, Schnase JL. MERRA/Max: A Down-Selection and Monte Carlo Convergence-Based Modeling of Complex Systems. AGU Fall Meeting Abstracts. San Francisco: AGU; 2019. Available: https://agu.confex.com/agu/fm19/meetingapp.cgi/Paper/496296

161. Stahle DW. Anthropogenic megadrought. Science. 2020;368: 238–239. doi:10.1126/science.abb6902

162. Williams AP, Cook ER, Smerdon JE, Cook BI, Abatzoglou JT, Bolles K, et al. Large contribution from anthropogenic warming to an emerging North American megadrought. Science. 2020;368: 314–318. doi:10.1126/science.aaz9600

163. The North American Monsoon | NOAA Climate.gov. 2021 [cited 29 Jun 2023]. Available: http://www.climate.gov/news-features/blogs/enso/north-american-monsoon

164. Zeng J, Zhang Q. The trends in land surface heat fluxes over global monsoon domains and their responses to monsoon and precipitation. Sci Rep. 2020;10: 5762. doi:10.1038/s41598-020-62467-0

165. NOAA. The North American Monsoon. NOAA NWS Climate Prediction Center; 2019 p. 25. Available: https://www.cpc.ncep.noaa.gov/products/outreach/Report-to-the-Nation-Monsoon_aug04.pdf

166. Adams DK, Comrie AC. The North American Monsoon. Bull Amer Meteor Soc. 1997;78: 2197–2213. doi:10.1175/1520-0477(1997)078<2197:TNAM>2.0.CO;2

167. Liebmann B. Characteristics of North American Summertime Rainfall with Emphasis on the Monsoon. American Meteorological Society Jounal of Climate. 2008;21: 1277–1294.

168. Adams DK. Review of Variability in the North American Monsoon. 2016 [cited 1 Jul 2023]. Available: https://geochange.er.usgs.gov/sw/changes/natural/monsoon/

169. Hobeichi S, Abramowitz G, Ukkola AM, De Kauwe M, Pitman A, Evans JP, et al. Reconciling historical changes in the hydrological cycle over land. npj Clim Atmos Sci. 2022;5: 17. doi:10.1038/s41612-022-00240-y

170. Hamal K, Sharma S, Khadka N, Baniya B, Ali M, Shrestha MS, et al. Evaluation of MERRA-2 Precipitation Products Using Gauge Observation in Nepal. Hydrology. 2020;7: 40. doi:10.3390/hydrology7030040

171. Lehmann P, Merlin O, Gentine P, Or D. Soil Texture Effects on Surface Resistance to Bare-Soil Evaporation. Geophysical Research Letters. 2018;45. doi:10.1029/2018GL078803

172. Lian X, Zhao W, Gentine P. Recent global decline in rainfall interception loss due to altered rainfall regimes. Nat Commun. 2022;13: 7642. doi:10.1038/s41467-022-35414-y

173. Cheng Y, Chan PW, Wei X, Hu Z, Kuang Z, McColl KA. Soil Moisture Control of Precipitation Re-evaporation over a Heterogeneous Land Surface. Journal of the Atmospheric Sciences. 2021;78: 3369–3383. doi:10.1175/JAS-D-21-0059.1

174. Lortie CJ, Filazzola A, Westphal M, Butterfield HS. Foundation plant species provide resilience and microclimatic heterogeneity in drylands. Sci Rep. 2022;12: 18005. doi:10.1038/s41598-022-22579-1

175. Varner J, Dearing MD. The importance of biologically relevant microclimates in habitat suitability assessments. Moreau CS, editor. PLoS ONE. 2014;9: e104648. doi:10.1371/journal.pone.0104648

176. Porter WP, Budaraju S, Stewart WE, Ramankutty N. Calculating Climate Effects on Birds and Mammals: Impacts on Biodiversity, Conservation, Population Parameters, and Global Community Structure. American Zoologist. 2000;40: 597–630. doi:10.1668/0003-1569(2000)040[0597:cceoba]2.0.co;2

177. Ruth JM. Cassin’s Sparrow Status Assessment and Conservation Plan. Biological Technical Publication BTP-R6002-2000. Denver, CO: U.S. Department of the Interior, Fish and Wildlife Service; 2000.

178. Lynn J. Cassin’s Sparrow (Aimophila cassinii): A Technical Conservation Assessment. USDA Forest Service, Species Conservation Project. Rocky Mountain Region. 2006;46.

179. Anderson JT, Conway WC. The flight song display of the Cassin’s Sparrow (Aimophila cassinii): form and possible function. Bulletin of the Texas Ornithological Society. 2000;33: 1–12.

180. New Database available: USGS Releases “Species of Greatest Conservation Need” Lists | U.S. Geological Survey. [cited 3 Jul 2023]. Available: https://www.usgs.gov/news/technical-announcement/new-database-available-usgs-releases-species-greatest-conservation-need

181. Programs: Fish and Wildlife: Threatened and Endangered: State T&E Data: New Mexico | Bureau of Land Management. [cited 3 Jul 2023]. Available: https://www.blm.gov/programs/fish-and-wildlife/threatened-and-endangered/state-te-data/new-mexico

182. NASA Case Study – Amazon Web Services (AWS). In: Amazon Web Services, Inc. [Internet]. 2021 [cited 26 May 2021]. Available: https://aws.amazon.com/partners/success/nasa-image-library/

183. Research and Technical Computing on Amazon Web Services (AWS). In: Amazon Web Services, Inc. [Internet]. 2021 [cited 26 May 2021]. Available: https://aws.amazon.com/government-education/research-and-technical-computing/

184. Google Cloud offers global support for academic research. In: Google [Internet]. 2019 [cited 26 May 2021]. Available: https://blog.google/products/google-cloud/google-cloud-offers-global-support-for-academic-research/

185. Our head’s in the cloud, but we’re keeping the earth in mind. In: Google Cloud Blog [Internet]. 2019 [cited 26 May 2021]. Available: https://cloud.google.com/blog/topics/google-cloud-next/our-heads-in-the-cloud-but-were-keeping-the-earth-in-mind/

186. Cloud Computing Services | Microsoft Azure. 2021 [cited 26 May 2021]. Available: https://azure.microsoft.com/en-us/

187. Data Science Virtual Machines | Microsoft Azure. 2021 [cited 26 May 2021]. Available: https://azure.microsoft.com/en-us/services/virtual-machines/data-science-virtual-machines/

188. Copernicus Climate Data Store (CDS). 2021 [cited 8 Nov 2021]. Available: https://cds.climate.copernicus.eu/#!/home

189. Copernicus Climate Change Service. Global bioclimatic indicators from 1950 to 2100 derived from climate projections. ECMWF; 2021. doi:10.24381/CDS.A37FECB7

190. Copernicus Climate Change Service. Downscaled bioclimatic indicators for selected regions from 1979 to 2018 derived from reanalysis. ECMWF; 2021. doi:10.24381/CDS.FE90A594

191. About Copernicus | Copernicus. [cited 5 Jul 2023]. Available: https://www.copernicus.eu/en/about-copernicus

192. Rinnan DS, Lawler J. Climate-niche factor analysis: a spatial approach to quantifying species vulnerability to climate change. Ecography. 2019;42: 1494–1503. doi:10.1111/ecog.03937

193. Foden WB, Young BE, Akçakaya HR, Garcia RA, Hoffmann AA, Stein BA, et al. Climate change vulnerability assessment of species. Wiley interdisciplinary reviews: climate change. 2019;10: e551.

194. Sang Z, Hamann A. Climatic limiting factors of North American ecosystems: a remote-sensing based vulnerability analysis. Environ Res Lett. 2022;17: 094011. doi:10.1088/1748-9326/ac8608

195. Hatten JR, Giermakowski JT, Holmes JA, Nowak EM, Johnson MJ, Ironside KE, et al. Identifying bird and reptile vulnerabilities to climate change in the Southwestern United States. US Geological Survey; 2016.

196. Peterson AT, Papeş M, Soberón J. Mechanistic and correlative models of ecological niches. European Journal of Ecology. 2015;1: 28–38. doi:10.1515/eje-2015-0014

197. Dormann CF, Schymanski SJ, Cabral J, Chuine I, Graham C, Hartig F, et al. Correlation and process in species distribution models: bridging a dichotomy: Bridging the correlation-process dichotomy. J Biogeogr. 2012;39: 2119–2131. doi:10.1111/j.1365-2699.2011.02659.x

198. Enriquez-Urzelai U, Kearney MR, Nicieza AG, Tingley R. Integrating mechanistic and correlative niche models to unravel range-limiting processes in a temperate amphibian. Global Change Biology. 2019;25: 2633–2647. doi:10.1111/gcb.14673

199. Evans TG, Diamond SE, Kelly MW. Mechanistic species distribution modelling as a link between physiology and conservation. Conserv Physiol. 2015;3: cov056. doi:10.1093/conphys/cov056

200. Buckley LB, Urban MC, Angilletta MJ, Crozier LG, Rissler LJ, Sears MW. Can mechanism inform species’ distribution models?: Correlative and mechanistic range models. Ecology Letters. 2010; no-no. doi:10.1111/j.1461-0248.2010.01479.x

201. Van der Meersch V, Chuine I. Estimating process-based model parameters from species distribution data using the evolutionary algorithm CMA-ES. Methods Ecol Evol. 2023;14: 1808–1820. doi:10.1111/2041-210X.14119

202. Schnase JL, Carrol ML. The MMX Toolkit: High performance, reanalysis-based climatic suitability modeling to advance avian conservation. Proceedings of the 2023 Conference on Big Data from Space (BiDS’23). Vienna, Austria; 2023.

